# Analysis of phloem trajectory links tissue maturation to cell specialization

**DOI:** 10.1101/2021.01.18.427084

**Authors:** Pawel Roszak, Jung-ok Heo, Bernhard Blob, Koichi Toyokura, Maria Angels de Luis Balaguer, Winnie W. Y. Lau, Fiona Hamey, Jacopo Cirrone, Xin Wang, Robertas Ursache, Hugo Tavares, Kevin Verstaen, Jos Wendrich, Charles W. Melnyk, Dennis Shasha, Sebastian E. Ahnert, Yvan Saeys, Bert De Rybel, Renze Heidstra, Ben Scheres, Ari Pekka Mähönen, Berthold Göttgens, Rosangela Sozzani, Kenneth D. Birnbaum, Yrjö Helariutta

## Abstract

The mechanisms that allow cells in the plant meristem to coordinate tissue-wide maturation gradients with specialized cell networks are critical for indeterminate growth. Here, we reconstructed the protophloem developmental trajectory of 19 cells from cell birth to terminal differentiation at single cell resolution in the Arabidopsis root. We found that cellular specification is mediated near the stem cell niche by PHLOEM EARLY DOF (PEAR) transcription factors. However, the PEAR dependent differentiation program is repressed by a broad gradient of PLETHORA (PLT) transcription factors, which directly inhibit PEARs’ own direct target *ALTERED PHLOEM DEVELOPMENT (APL)*. The dissipation of PLT gradient around 7 cells from the stem cell activates APL expression, and a subsequent transitional network that results in a “seesaw” pattern of mutual inhibition over developmental time. Together, we provide a mechanistic understanding of how morphogen-like maturation gradients interface with cell-type specific transcriptional regulators to stage cellular differentiation.

## Main text

Roots consist of several concentric layers of functionally distinct cell files, which initially bifurcate and establish distinct identities around the quiescent center and its surrounding stem cells. Cells within each file mature through the distinct zones of cell proliferation and differentiation (*1*). For example, in Arabidopsis the development of the protophloem sieve elements (PSEs) involves a transient period of cell proliferation, during which, in addition to amplification of cells within the file, two lineage-bifurcating events take place (Fig. 1A) (*2*). Soon after the cell proliferation ceases, cells of the PSE lineage initiate a differentiation process which culminates in enucleation, an irreversible process that gives rise to the mature conductive cells (*3*). Because of specific modulation of the graded distribution of the key phytohormonal cue auxin, the differentiation of PSEs occurs faster than that of the other cell files (*4*). Therefore, PSE development offers a traceable scheme to understand how the two processes of cell specialization and maturation interact.

**Figure 1.**
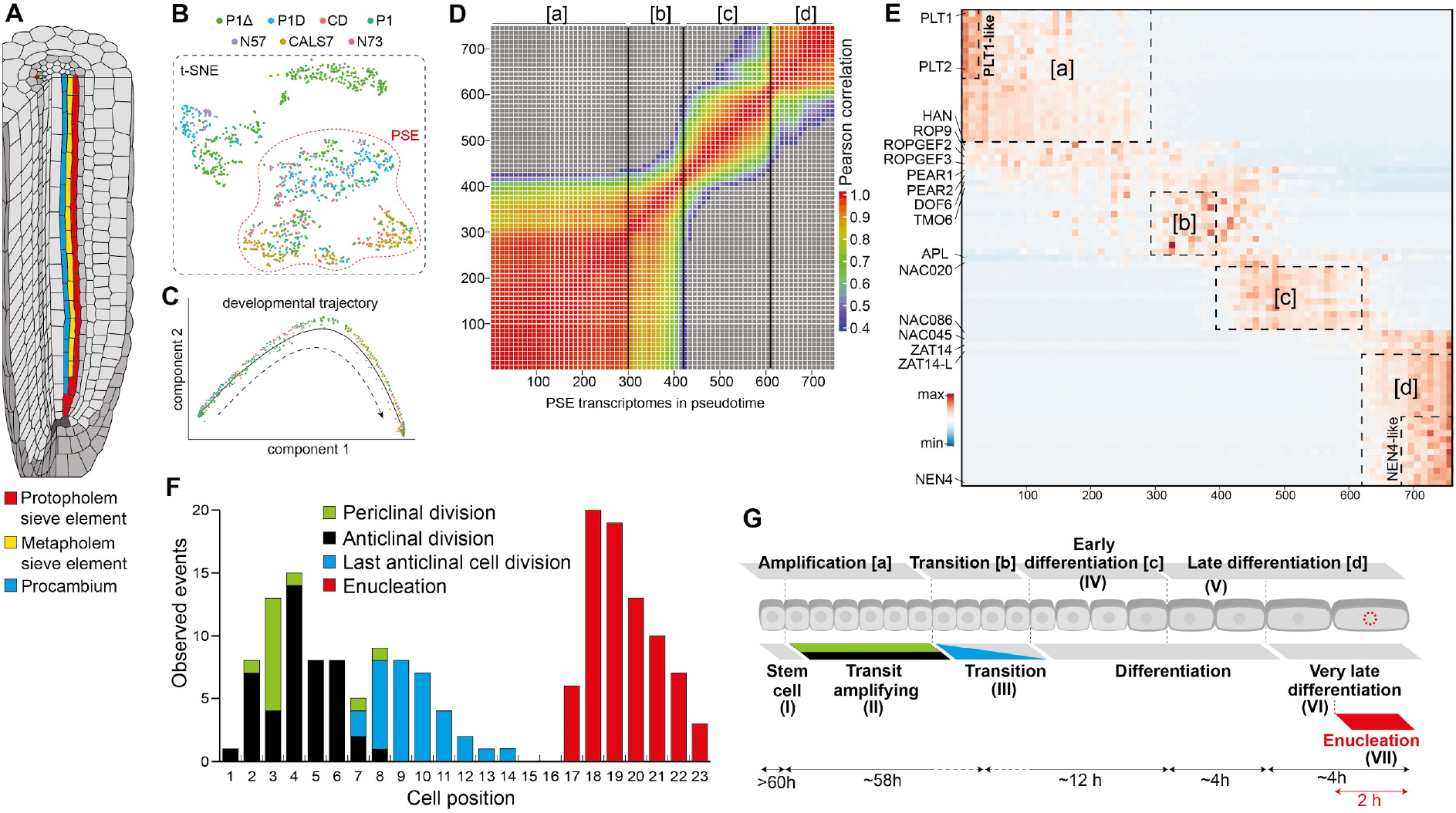
Phloem development at single-cell resolution. A) Schematic of the Arabidopsis root tip depicting position of PSE, MSE and procambium cell lineages originating from a single phloem stem cell. B) t-SNE plot of 1242 transcriptomes of cells sorted with P1Δ, P1D, CD, P1, N57, CALS7 and N73 reporter lines specific to different domains of the developing phloem. Indicated PSE cells were used for the pseudotime trajectory analysis (fig. S2, Supp Info). C) PSE transcriptomes ordered along developmental trajectory using Monocle 2. D) Heatmap of Pearson correlation along the pseudotime trajectory. Vertical lines indicate 3 strongest correlation drops and separate four groups of transcriptomes with higher similarity [a], [b], [c] and [d]. E) Gene expression heatmap of important PSE regulators and 10 most specific genes from the 4 groups defined in D) and the nested *PLT1* (“PLT1-like”) or *NEN4* (“NEN4-like”) expression domains in pseudotime-ordered PSE transcriptomes. F) Histogram of cell behaviour based on long-term live imaging. G) Seven domains and the time cells spend in each position of the developing PSE as determined by the transcriptomics (above) and live imaging (below): (I) “stem cell”, position 1 [a], t>60h; (II) “transit amplifying”, position 2-9 [a], t=58h, SD+8.1h, (III) “transitioning”, position 8-11 [b]; (IV) “early differentiating”, position 10-15 [c], t=12h; (V) “late differentiating”, position 16-17 [d], t=4h; (VI) “very late differentiating - NEN4-like”, position 18-19 [d], t=4h; VII “enucleating”, position 19 [d], t= 2h (Movie S1, S2).

In order to understand the process of PSE development at a high resolution, we took a combination of approaches based on live-imaging (*5*) and single cell transcriptomics (*6*). Using time-lapse confocal imaging with a phloem-specific marker (*pPEAR1::H2B-YFP pCALS7::H2B-YFP*) we precisely mapped cellular behavior of the (on average) 19 cells that constitute the PSE developmental trajectory. The passage of the cell from its “birth” at the stem cell until its enucleation took a minimum of 79 h (fig. S1, movies S1, S2). To dissect the genetic control underlying this temporal progression, we performed fluorescence-activated cell sorting (FACS) in order to isolate cells for single-cell transcriptomics (*6*–*12*) with fluorescent reporter lines whose expression represent various spatio-temporal domains within the PSE trajectory (fig. S2A, B), as well as with a modified *pPEAR1Δ::erVenus* reporter line that represents the bifurcating procambial and metaphloem trajectories in addition to the PSE trajectory (fig. S2A). The single cell profiles allowed us to use T-Distributed Stochastic Neighbor Embedding together with known PSE markers to identify 758 cells that densely sampled the 19 cell positions that captured PSE maturation (Fig. 1B, fig. S2C-G).

We sought to use the high-resolution profile of the PSE lineage to ask how cell passage through stable signaling gradients in the meristem controls the stages of cellular specialization. In particular, while a number of regulators of either phloem cell identity or meristem maturation have been described (*13*, *14*), there is little known about how these two important regulatory processes interact to control organogenesis. Using Monocle 2 (*15*, *16*) we projected the 758 PSE cells into a pseudo-temporal order and investigated transcriptional transitions along the developmental trajectory (Fig. 1B-D). Rather than gradual changes, we observed three dramatic shifts in transcriptomic patterns (Fig. 1D, E; Table S2). Based on the alignment of these four transcriptome domains to the temporal expression patterns of selected genes, we were able to determine that these domains correspond approximately to cells at positions 1-7 [a], 8-11 [b], 12-15 [c] and 16-19 [d], respectively (fig. S3). To further understand which aspects of PSE maturation these various positions represent, we extended the time-lapse confocal imaging with more temporally specific marker lines *pNAC86::H2B-YFP* and *pNEN4::H2B-YFP*. We found that the differentiation time, measured from the last cell division till enucleation, is variable up to the final stage defined by *NAC45/86-DEPENDENT EXONUCLEASE-DOMAIN PROTEIN 4* (*NEN4*) expression (active in positions 18-19), Fig. 1E, fig. S1D, H, I, Movies S1-S12). In summary, based on the high congruence of the single-cell transcriptome and live imaging data, we were able to assign seven distinct cell behavior domains along the PSE trajectory (Fig. 1F, G, fig. S1).

The first major transition in the PSE cellular trajectory corresponded to a shift in cellular behavior from transit amplifying cells to transitioning cells that mapped closely to the first major change in the PSE transcriptome. In the first transcriptional zone, we detected PLETHORA (PLT) transcripts (Fig. 1E), whose relatively persistent proteins are known to spread shootward through cell-cell movement and to form a gradient through mitotic dilution (*14*). The PLTs promote cell division at moderate concentrations, and as the PLT levels drop, cells initiate elongation and differentiation (*14*, *17*, *18*). However, it is not clear how individual cell files interpret the PLT gradient for their own specialized differentiation.

We hypothesized that the PLT gradient might mediate the first major transition (i.e. domain [a] to [b]) towards PSE differentiation by permitting a new set of transcripts to be expressed (Fig. 2A). We tested this hypothesis by driving PLT2 under several promoters that extended its expression in the PSE in later maturation stages than its native domain (Fig. 2B, fig. S4A). When using the *pNAC86::XVE* inducible promoter (*3*, *19*), ectopic PLT2 delayed PSE enucleation (Fig. 2B, fig. S4A) and transcriptional profiling of phloem cells expressing the construct showed an upregulation of genes (Table S3) that mapped to early stages of the PSE single-cell trajectory (from domains I-II) - the known PLT2 protein gradient (Fig. 2C). These results suggest that extending the PLT2 gradient is sufficient to prolong the early stages of meristem maturation within the PSE lineage, providing a connection between cellular maturation and meristem-wide protein gradient. In addition, in the pseudo-time ordered single cells, we could detect complementary oscillatory patterns of the putative S-phase and G2-M-phase genes that were upregulated PLT targets, apparently corresponding to regular progressions through the cell-cycle (Fig. 2C, fig. S4B). Furthermore, *ALTERED PHLOEM DEVELOPMENT* (*APL*), *NAC45/86* and *NEN4*, known key regulators of the PSE enucleation pathway (*3*), were among the PLT2-downregulated genes (table S3). This is consistent with the presence of *APL* in the large set of genes downregulated by PLT overexpression (*18*). We validated the downregulation of *APL* and *NEN4* by ectopic PLT2 expression with in situ hybridization (Fig. 2D, fig. S4C). We also temporally monitored a shootward shift of *APL* expression domain in the roots expressing PLT2 in the phloem cells beyond its native domain confirming that activation of APL-dependent genetic program requires dissipation of the PLT gradient (Fig. 2E). In order to test the requirement of PLTs in maintaining transit amplifying phloem cell pool, we used an inducible, tissue specific CRISPR/Cas9 approach to mutate *PLT2* specifically in PSE cell file (*20*). We observed a shift of the PSE differentiation as well as the expression of *pAPL::erTurq* reporter towards the QC without affecting meristem size or root growth, showing that loss of PLT function in its native domain allows precocious expression of mid-to late-stage PSE differentiation regulators (Fig. 2F, fig. S4E-G).

**Figure 2.**
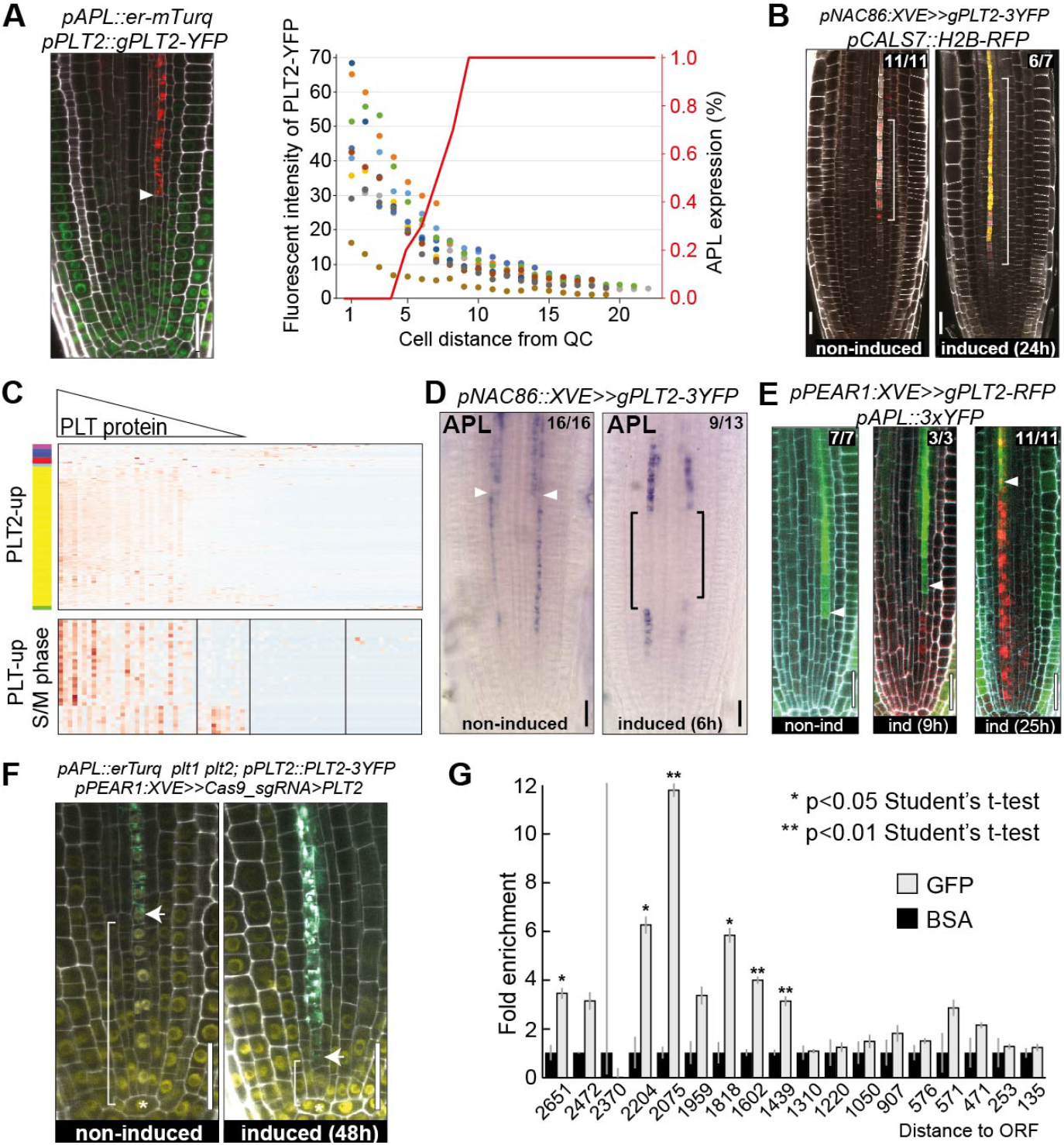
PLT2 inhibits phloem differentiation by directly repressing APL expression. A) Quantification of fluorescent intensity of PLT2-YFP in PSE cells of 9 roots indicated with dots of different colours. Percentage of roots expressing *APL* in a given PSE cell is indicated as a red line (n=9). Onset of *APL* expression coincides with diminishing level of PLT2 protein. Arrowhead indicates onset of *APL* expression in PSE. B) Ectopic expression of *PLT2* under *pNAC86::XVE* promoter delays PSE enucleation. Square brackets indicate extended expression domain of *pCALS7::H2B-RFP*, a reporter used for monitoring enucleation. C) Native expression profile of PLT2 targets in PSE cells ordered in pseudotime. Genes upregulated after 6 hours of induction of the line shown in B) are plotted. Upper panel shows gradually diminishing expression of target genes which reflects the PLT2 protein gradient. Lower panel shows PLT2 upregulated cell cycle genes with oscillatory expression pattern. D) In situ hybridization of *APL* before and 6h after ectopic expression of PLT2-3xYFP. Arrowheads indicate position of PSE enucleation beyond which point *APL* is expressed in phloem pole pericycle, companion cells and metaphloem sieve element (fig. S4D). Brackets indicate *pNAC086* activity domain. E) Time course of transcriptional repression of *APL* in cells ectopically expressing PLT2-RFP under inducible *pPEAR1::XVE* promoter. F) Early activation of *APL* expression 48h after phloem specific knock-out of *PLT2*. G) ChIP-qPCR of PLT2-3xYFP on *APL* promoter revealed PLT binding region −2204 to −1439 bp upstream of *APL* ORF. All scale bars, 25 μm.

We sought to further test whether PLT2 directly regulates the PSE-specific differentiation program, as we found AP2 (PLT2 family) family binding sites in the *APL* promoter region (*21*). Indeed, we confirmed the direct binding of PLT2 to several regions of the *APL* promoter by ChIP-qPCR (Fig. 2G). Furthermore, along with AP2 sites, the *APL* promoter is also enriched for HANABA TANARU (HAN), a GATA transcription factor, binding sites. In turn, *HAN* is a PLT target (*18*) and accordingly, upon ectopic PLT2 expression we can detect *HAN* transcripts during late PSE development (fig. S4H). Ectopic HAN expression under *pNAC86:XVE* led to a delay in enucleation (fig. S4I), similar to PLT2 overexpression in the same domain. We conclude that the PLT gradient directly (and possibly in a feedforward manner with HAN) orchestrates PSE differentiation by cell autonomously repressing transcription of the phloem regulator *APL*. Overall, the results show how the PLT gradient first orchestrates cell proliferation in the PSE lineage and then helps to time the later stages of cellular maturation.

Given the results above, we reasoned that an early phloem-specific transcription factor must activate *APL* expression. In order to identify phloem enriched genes, we utilized RNAseq data of bulk-sorted PSE cells and a previously published tissue specific data set (*22*) (table S4, fig. S5A). Subsequently, these were analyzed for sieve element enrichment using a force directed approach on the 272 single cell transcriptomes originating from the *pPEAR1Δ::erVenus* reporter line (Fig. 3A, B, fig. S5B-H, table S5). We identified 542 genes (table S6) and examined their specificity in the published whole-root scRNAseq atlas (table S7)(*12*). Furthermore, we modeled gene regulation using a machine learning approach on the pseudotime-ordered 758 single-cell profiles and 4924 highly variable genes. Among 208 TFs in this dataset, the majority of known PSE transcription factors (such as *APL*, *NAC045* and *NAC086*) were among the top 20 regulators (table S8). We validated the model by comparing predicted targets with genes induced by *in vivo* ectopic expression of the same TFs, confirming a significant overlap of targets in 3 out of 5 cases (table S8). Among the top 20 regulators we also identified four related genes that encode early SE abundant PEAR transcription factors (Fig. 3C). The simultaneous loss of six PEAR genes was recently shown to result in defects in PSE differentiation (*23*). We subsequently profiled the transcriptomes of wildtype and pear sextuple mutant (Fig. 3D) root meristems and identified 203 downregulated genes overlapping with our PSE specific gene list (table S9). Noticeably, the expression of *APL* as well as its downstream targets – *NAC086* and *NEN4* was strongly reduced in the pear sextuple mutant (Fig. 3E, fig. S6A). Consistently with our RNA-seq data, a fluorescent reporter of *pAPL* did not show any expression in the PSE in the pear sextuple mutant but its expression was restored upon induction of *PEAR1*, placing *APL* downstream of PEAR regulation (Fig. 3F).

**Figure 3.**
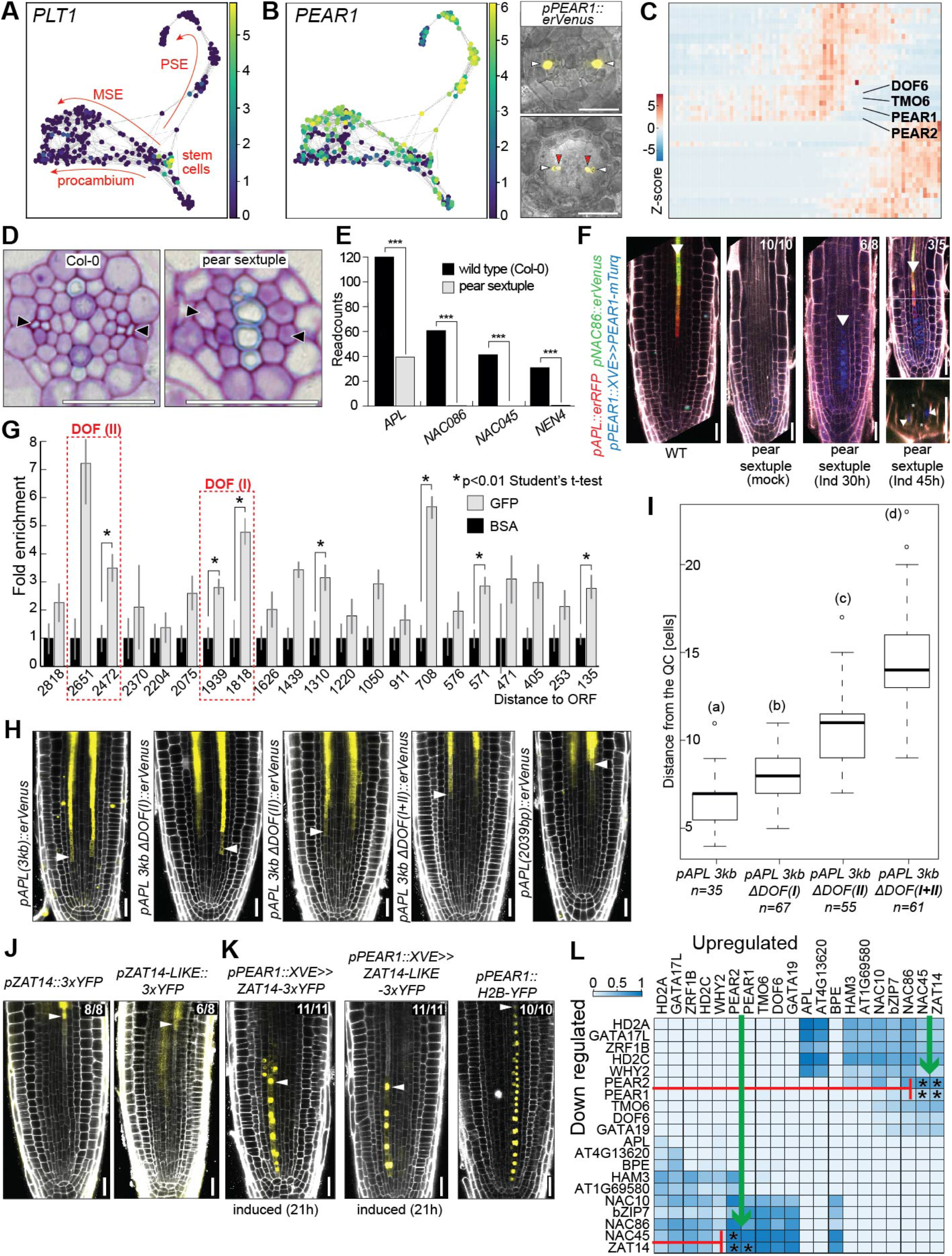
PEARs are important regulators of phloem differentiation. A) Force-directed graph of 272 single-cell transcriptomes obtained using the *pPEAR1Δ::erVENUS* reporter. Plotted is expression of stem cell abundant *PLT1.* Arrows: cellular trajectories inferred from known gene expression patterns (fig. S5). B) Strong enrichment of *PEAR1* expression in PSE and MSE trajectories confirmed by *pPEAR1::erVENUS* reporter line. White arrowheads: PSE, red arrowheads: MSE. C) Expression heatmap: PEAR genes among the earliest phloem specific transcription factors. D) Lack of PSE differentiation in the mature part of the pear sextuple mutant root. Arrowheads: PSE position E) Lack of APL pathway activation in the roots of pear sextuple mutant based on RNASeq analysis. F) Inducible expression of *PEAR1-mTurq* is sufficient to activate transcription of *pAPL* and *pNAC86* reporters in pear sextuple background. G) ChIP-qPCR of PEAR1-YFP shows direct interaction of PEAR1 with *APL* promoter at multiple positions. Two prominent PEAR1 binding sites are indicated with red dashed rectangles. H) Expression patterns of modified *pAPL* reporter lines. Length of “3kb” promoter equals 2962 bp. DOF(I) and DOF(II) correspond to two enhancer elements indicated in panel G. Details of modification are provided in fig. S6C. I) Quantification of onset of *pAPL* expression after modification of DOF binding motives. Statistically significant differences between groups were tested using Tukey’s HSD test *P* < 0.05. Different letters indicate significant difference at *P* < 0.05. J) Expression of ZAT14 and ZAT14L during late differentiation of PSE. Arrowheads: last cell before enucleation. K) Ectopic expression of *ZAT14* and *ZAT14L* under *pPEAR1* results in cell elongation and inhibition of cell division. Arrowheads: last cell before enucleation. *pPEAR1::H2B-YFP* line shows regular number of PSE cells. L) Heatmap shows significantly overlapping and oppositely regulated target sets of the 20 most important TFs from the GRN model. Color intensity shows a fraction of overlapping target sets. The colormap represents significantly overlapping sets (Fisher Exact Test, if p<0.05, val=1) multiplied by the fraction of overlap. Asterisk indicates experimental validation of up and downregulated sets from TF OE *in vivo* (tables S15, S16). All scale bars, 25 μm.

To test whether PEAR1 can directly regulate expression of *APL*, we performed chromatin immunoprecipitation (ChIP) followed by quantitative PCR (qPCR) using *pPEAR1::PEAR1-GFP* protein fusion and identified multiple PEAR1 binding sites within *APL* promoter (Fig. 3G). Truncation analysis of *pAPL* indicated presence of an enhancer element, responsible for expression of *APL* in the cells transitioning from cell division to cell differentiation, within 2039 bp to 2962 bp region upstream of *APL* open reading frame (ORF) (Fig. 3H). Our ChIP analysis detected a single strong PEAR1-GFP peak in the promoter sequence beyond 2039 bp distance from the ORF and another strong peak at the end of 2 kb region, both of which were also detected in the publicly available DAP-Seq data (Fig. 3G, fig. S6C)(21).

Furthermore, within the detected regions (−2672 to −2512 and −1946 to −1844) we identified multiple clusters of DOF binding motifs (AAAG)(21) that constitute an enhancer element required for the transcriptional activation of *APL* in the phloem transition zone (domain III) (Fig. 3H, I, fig. S6C). Collectively, our data supports a central role of a PEAR-APL causal relationship in the genetic pathway controlling PSE differentiation. In addition, PLT factors mediate transit amplifying divisions that, in part, lead to the dissipation of their own gradient. This, in turn, relieves the repression of phloem-specific PEAR target *APL*, thus timing onset of differentiation. These opposing effects on *APL* regulation illustrate the antagonism between regulators that stage maturation and differentiation.

The final major transcriptional transition occurs between the domains IV-V. To further explore this transition, we ectopically expressed NEN4 and PLT2 at various developmental stages. Whereas in the earlier domains ectopic NEN4 expression causes cell death and PLT expression renders the cells back to the cell cycle, the developmental program of domain V appears resilient to these perturbations (Fig. 2B, fig. S4A, fig. S8). This indicates that the high number of PSE specific genes during the final 8h of differentiation remodel the cellular behavior in an irreversible manner. We next sought to explore how widely the PEARs control transcriptional programs related to this final stage of sieve element development. We combined a gene regulatory analysis in the pear mutant with systematic overexpression and modelling approaches (fig. S6A, B, fig. S7). Our analysis revealed that - in addition to known phloem regulators *APL, NAC045, NAC086* and *NAC028* −10 out of 13 newly validated phloem enriched transcription factors are dependent on PEARs (fig. S6A, B, Fig. 3F). Overexpression of two of these, *ZAT14* (AT5G03510), which was also the 3rd most important TF in the machine learning model, and its close homolog *ZAT14L* (AT5G04390) led to arrest of cell cycle and premature cell elongation (Fig. 3J, K). Transcriptional profiling provided further evidence for putative dual role in timing cell division and cell expansion (that occurs largely after enucleation in this cell lineage) (tables S10-S14). In addition, the gene regulatory network model predicted a pattern of sequential mutual inhibition in the target sets of high-scoring transcriptional regulators (table S15); for example, genes repressed by ZAT14 significantly overlap with genes activated by the earlier expressed PEARs and *vice versa* (Fig. 3L). Overexpression analysis confirmed a significant over-representation in the overlap between genes up-regulated by PEARs and down-regulated by ZAT14 (table S16)(*23*).

Recent work has shown that high-throughput single-cell RNA-seq profiles can be used to assemble the anatomy of the Arabidopsis root (*6*–*12*). By combining single-cell transcriptomics with live imaging, here we have mapped the cellular events from the birth of the sieve element cell to its terminal differentiation spanning a timeframe of 80 h. In addition, the PEARs activate the 20-hour terminal differentiation program, which highlights them as central integrators that connect early and late phloem development. The late, PEAR-regulated PSE program is directly and antagonistically controlled by the broad PLT gradient, which connects this morphogen-like gradient to cellular maturation. Although the gradually diminishing PLT gradient stages meristem differentiation (*14*, *17*, *18*), our high-resolution phloem trajectory reveals three abrupt transitions in the gene expression program. We propose that mutual inhibition of target genes by sequentially expressed transcription factors represents a “seesaw” mechanism (fig. S9) that allows rapid transitions and prevent gene expression programs with conflicting effects on cellular physiology (e.g., division vs. enucleation). Similar models have been implicated in so-called attractor states in cell fate decisions in animals (*24*). In the future it will be interesting to determine how conserved these principles of sieve element differentiation are in an evolutionary context, as well as how extensively they apply to the other differentiation trajectories in plants.

## Materials and Methods

### Plant materials and growth condition

Previously published and newly generated plant materials are listed in Table S1.

Arabidopsis seeds were surface sterilized in either a bleach solution (5% sodium hypochlorite with 0.02% tween 20) for 5 minutes with gentle rotation followed by 8 washing steps with sterile distilled water, or by chlorine gas treatment (3ml HCl added to 100ml of bleach enclosed in a airtight container) for a minimum of 4h. After sterilization seeds were imbibed in sterile water for 2 – 5 days at 4 °C in the dark. Seeds were germinated on 1/2 strength Murashige and Skoog (½ MS) agar medium containing 1% sucrose and 0.8% difco agar. Seedlings and adult plants were grown under long-day condition – 16 hours light (188 μmol m^−2^ s^−1^) and 8 hours dark at 23 °C. For confocal imaging, anatomical analysis, and RNA-seq experiments, 5-day-old vertically grown seedlings were used, unless otherwise specified.

### Generation of fluorescent reporter lines

All the reporter lines were generated according to the previously described method which is based on multisite gateway system (Invitrogen)(*3*). To generate transcriptional reporters, promoters were cloned into pDONR P4-P1R entry vector. In general, up to 3 kb upstream region of each gene was PCR amplified using Col-0 genomic DNA as a template and cloned by BP cloning method (Invitrogen). Inducible constructs for *pHAN*, *pNEN4*, and *pNAC73* were generated by a classical restriction/ligation approach in the *p1R4-ML::XVE* backbone vector using T4 ligase as described in (*19*). Constructs with small deletion (−1039 to −853) in the promoter sequence of *PEAR1* and modifications in DOF(I) fragment of *pAPL* (fig. S6C) were generated according to the NEBuilder HiFi DNA Assembly Master Mix protocol (New England Biolabs). Modified DOF(II) sequence of *pAPL* was synthetized by Twist Bioscience, USA, and cloned into entry clone with *pAPL* following NEBuilder protocol. Genomic fragments for *GIS*, *AT2G45120*, *NAC010*, *NTL8*, *BPEp2*, *NAC048*, *NAC075*, *WRKY21*, ZAT14 (AT5G03510) and *ZAT14L* (AT5G04390) were cloned into pDONR221. Histone 2B (H2B), DBOX domain from *A. thaliana* Cyclin B1;1 or fluorescent proteins were used in the second entry vector, followed by a fluorescent protein or terminator in the third entry vector, respectively. CRISPR knock-out construct for PLT2 in pDONR221 is a courtesy of Ari-Pekka Mahonen’s group (*20*). Primer sequences are listed in Table S19. Agrobacterium-mediated floral dip method was used to generate transgenic Arabidopsis plants.

### Protoplast isolation and cell sorting

Arabidopsis root protoplasts were isolated according to the published protocol (*25*). In short, sterilized seeds were plated onto the nylon mesh (57-103, Nitex) placed on the top of MS agar media. Two rows of seeds comprising approximately 100-150 seeds per row were then grown vertically for 5 days in long-day growth condition. To isolate root protoplasts, ⅓ of the root was cut and chopped on the surface of the mesh and transferred to the buffer containing cell wall degrading enzymes −1.5 g Cellulase (C1794, Sigma) and 0.1 g Pectolyase (P3026, Sigma) in 100 ml protoplasting solution. After 1 hour of incubation with occasional stirring, the protoplasts were spun down (200 rcf for 6 min) in 15 ml Falcon tube and after removal of supernatant, cells were resuspended in 700 μl protoplasting solution without enzymes. After filtering with 70 μm and 40 μm cell strainers sequentially, FACS was performed using 70 micron nozzle at 10 psi. Fluorescence positive cells were collected in a round-bottom polystylene tube that contains 150 - 200 μl RLT buffer with beta-mercaptoethanol (10 μl / 1 ml RLT buffer, RNeasy micro kit protocol, Qiagen). Cell sorting was performed for about 15 min and collected cells were immediately frozen on dry ice and kept in −80 °C for up to one week. For single cell transcriptomics, protoplasts were produced and isolated as described here, but sorted individually into 96-well plates. The only exception are the cells isolated with the *pPEAR1Δ::erVenus* reporter line. To enrich in meristematic cells, whole roots were submerged in 1.5 ml tubes of 0.5 ml cell wall degrading buffer. The roots were picked up and shaken with forceps every 2 minutes for 12 minutes. This proved to be sufficient for the enzymes to degrade the cell wall in the transition zone of the root. The shaking helped to separate the meristems from the remains of root. The remains of the root were then removed with forceps and the contents of the tubes were united and incubated for another 50 min following the same steps as for the other samples.

### RNA sequencing of FACS isolated cells

For bulk sorted samples, including sorted cells ectopically expressing PLT2.

Frozen cells were quickly thawed by adding the same volume of EtOH (150 - 200 μl), and RNA extraction was done using RNeasy Micro Kit (Qiagen) as previously described by (*25*). RNA integrity was measured with Agilent HS RNA TapeStation system following the manufacturer’s instructions. Samples of RNA integrity value (RIN) 6.3 and above were taken to the following cDNA synthesis step. Clontech Smarter Ultra Low Input Library Kit V4 was used for cDNA synthesis and amplification (18 PCR cycles). Samples were then submitted to Novogene for library construction and RNA sequencing on an Illumina HiSeq SE50 run.

### Analysis of bulk RNA-seq data for the identification of phloem-abundant genes

Raw counts have been aligned and mapped according to a standardized Bowtie 2 suit using the TAIR10 genome. Feature Counts was used to obtain read counts and differential gene expression analysis was performed using edgeR. In order to find phloem-abundant genes (925, fig. S5A, table S4), we’ve adopted Shannon-entropy based selection of tissue specific genes as described by (*28*). As a comparison dataset, we used published root map RNA-seq data of other tissue types (*22*).

### Smart-seq Single-cell RNA sequencing

Single protoplast cells were sorted into lysis buffer and sample preparation for RNA-sequencing was performed as previously described (*27*, *28*). Briefly, cDNA was reverse transcribed with SuperScript II Reverse Transcriptase (Invitrogen) and amplified with 23 PCR cycles using KAPA HiFi HotStart polymerase (Roche). Amplified PCR products were purified with Ampure XP Beads (Agencourt) at a volume ratio of 1:0.6 DNA:beads, and quantified using Quant-iT™ PicoGreen™ dsDNA Assay Kit. Libraries were prepared using Illumina Nextera XT DNA preparation kit and either Nextera XT index kit – 96 indexes or Nextera XT index kit v2. Pooled libraries were sequenced on the Illumina HiSeq4000 (single-end reads, 50-bp).

### Identification of PSE specific genes

Differentially expressed genes between specific pairs of clusters were calculated using a t-test with overestimated variance and Benjamini-Hochberg correction for adjusted p-values as implemented in the scanpy (*29*) rank_genes_groups function. All genes significantly enriched (q-value <0.05) in PSE cluster 8 or 9 or 10 or 12 when compared to clusters 3 and 6 and 11 formed a list of 1192 genes enriched in SEs in this data set (Table S5). By cross referencing these to the 925 phloem enriched genes, 542 sieve element specific genes were identified (fig S5H, Table S6).

### Single-cell data analysis

The transcriptome raw data of individual cells were mapped using HISAT2 (*30*), SAM files converted to BAM files using SAMTools (*31*). Subsequently Cuffquant and Cuffnorm were used to obtain read counts and FPKM values (*32*). These were then analysed using Monocle2 (*15*, *16*). We filtered out data of cells with <10.000 reads mapped to a unique locus, <20% reads mapped to TAIR10 genome and <2000 genes showing an expression above 0.1 (1242 of 1375 transcriptome). Only genes with expression in at least 50 transcriptomes passing these thresholds were considered in the further downstream analysis. Using Monocle2’s tSNE and clustering function, among 14 clusters, we removed 6 clusters (2, 3, 5, 7, 11, 12; Fig S2) that do not represent PSE cells (fig. S2). Columella cells were sorted with *pPEAR1Δ::erVenus* reporter that shows strong ectopic fluorescence in those cells. Those cells were identified according to expression of *GLV5* and *GLV7* (*33*)(Fig S2, Clusters 2, 5, 11). These cells were then also removed from further analysis. The remaining transcriptomes (1026) were plotted after dimensional reduction using tSNE again. It revealed a cluster of mainly *pNAC057::2xGFP* and *pPEAR1::dBOX-3xYFP* originating transcriptomes (Cluster 7) as well as two clusters of *pPEAR1Δ::erVenus* cells that were separated from the other cells’s transcriptomes (fig. S2, Clusters 3, 11). Using an unsupervised approach by clustering these cells into 10 clusters and looking for differentially expressed genes (DEG) between these clusters, we created a developmental trajectory based on the top 4000 DEG. These resulted in a PSE trajectory with a single branching point where transcriptomes of the three outstanding clusters of the original t-SNE were located (Clusters3, 7, 12, Fig S2). We suspected that the *pNAC057::2xGFP* and *pPEAR1::dBOX-3xYFP* transcriptomes would likely originate from contaminating cells, as this cluster also showed expression of AT2G22850/bZIP6, whose reporter line (S17) is specifically expressed in the phloem pole pericycle. Based on the lineage branching analysis, original clusters 3 and 12 consisting mostly of cells originating from *pPEAR1Δ::erVenus* reporter were assigned MSE and procambium identity.

To further investigate the lineage branching, filtered *pPEAR1Δ::erVenus* transcriptomes were analysed in scanpy v1.4 (*29*). Genes detected in fewer than 10 cells were filtered out, and data were normalised using the scanpy normalize_per_cell function with default parameters. Normalised counts were log(x+1) transformed and highly variable genes identified with the scanpy highly_variable_genes function with parameters max_mean=8 and min_mean=0.5. Principal component analysis was calculated on scaled gene expression, and a force directed graph visualisation calculated using the scanpy draw_graph function with a k=10 nearest neighbours calculated on the top 50 principal components. Cells were clustered using Louvain clustering (*34*) on the k=10 nearest neighbor graph with resolution=2. Looking at the expression of genes known to be differentially expressed between the procambial and sieve element lineages (Fig S5) allowed the identification of the PSE, MSE and procambial cell clusters.

### Pseudotime inference for PSE trajectory

After removing the outstanding clusters containing the columella, lateral root cap, phloem pole pericycle, MSE and procambium cells in the Monocle2 analysis, the remaining 758 cells were placed on a linear developmental trajectory using unsupervised tSNE clustering into 10 clusters and DDRTree dimensional reduction on the top 4000 DEGs between clusters. This step assigned a pseudotime value to each cell and ordering the transcriptomes according to these values was confirmed by the observed expression patterns of known PSE expressed genes (Fig. 1E).

### Pearson correlation calculation

The previously obtained pseudo-time order of 758 protophloem cells was used to calculate the Pearson correlation along pseudo-time. We applied a moving average window to reduce the effect of noise to this analysis. We tested window sizes of 5, 10, 15, 20, 30, and 50 cells moved by 5, 10, 15, 20 or 30 cells. Averaging 30 cells and moving 10 cell positions per step gave enough simplification while keeping a good level of detail. The Pearson correlation between the averaged transcriptomes was then calculated for all expressed genes as well as the 4000 genes from the pseudo-time ordering. By plotting the correlation matrix in a heatmap, we could identify the points in pseudo-time where the transcriptome changes rapidly. These changes were more pronounced looking only at the 4000 ordering genes, as these would also be among the top DEG along pseudo-time. These rapid changes could also be explained by an underrepresentation of these cells in our data set. However, the distribution of cells in pseudo-time shows that these points of big transcriptome changes do not overlap with the end or beginning of sorted populations in pseudo-time (fig S2G). Thus an equal sampling before and after these changes can be assumed, making the underrepresentation unlikely.

After defining the 4 major domains on 30-averages, 10-cell averages in that range of 40 cells (2 adjacent 30-cell windows with step size 10) were used to define the borders more precisely, with a final step using 5-cell averages that were split in the middle.

### Domain specific gene identification

We manually determined the border between stem cell and the second cell in pseudotime by the first noticeable drop of *PLT1* expression at pseudotime position 27. The *NEN4* expression domain was determined by the increase of *NEN4* transcripts beyond pseudotime position 691. As these are nested within domains 1 and 4, respectively, only the definitions for domain 1-4 were used to identify domain specific genes. These domains were compared using the same approach as for the cluster comparisons in the force-directed analysis. The highest q-value (qval_max) for each gene in the three possible comparisons were used for thresholding. For PLT1-like and NEN4-like domains, comparisons to all 5 other domains were used, respectively. Only genes with qval_max<0.05 were considered significantly enriched (Table S2).

### RNA extraction and sequencing of whole meristem samples

For the transcriptome profiling of PEAR sextuple mutant (*pear1 pear2 dof6 tmo6 obp2 hca2*) and inducible overexpression lines of ZAT14 and ZAT14L, primary root meristems were dissected under stereomicroscope using conventional needles at 5 days after germination. To facilitate cutting, plants were transferred from the ½ MS 1% agar plates to ½ MS 5% agar plates in bulk of 10. Dissected meristems (in bulk of 10) were then transferred to the droplet of RNlater solution kept in the cap of the test tube at room temperature. Tubes were stored in the fridge until RNA extraction with RNeasy Plant Mini Kit (Qiagen). RNA extraction from 100 root tips of wild type Arabidopsis would typically yield 2.5μg of good quality (RIN>8.5) total RNA of which 2μg were submitted for RNA-Seq analysis on the HiSeq-PE150 platform at Novogene. Overexpression of ZAT14 and ZAT14L was induced by transferring *pWOL:XVE>>ZAT14-mTurq* and *pWOL:XVE>>ZAT14L-mTurq* plants onto the media plates supplemented with 5μM 17-b-estradiol for 10 or 14h. Plants transferred to media plates supplemented with DMSO we used as controls.

### Quantification of fluorescent reporter lines

For quantification, confocal microscopy image files were directly imported into the image analysis software FIJI (*37*). Cell walls were outlined using the polygon selection tool in the red channel of PI stained root images. In addition to the PSE cells, four large areas within the root but not covering any PSE cells were drawn and the detected fluorescence considered as background levels. The average fluorescence in the YFP channel was then measured and normalized to the background fluorescence. The presence of nuclei in images of *pCALS7::H2B-RFP*, *pCALS7::H2B-YFP* and *pNAC073::H2B-YFP* was judged in the corresponding channel. In case of *pCALS7::H2B-YFP* and *pNAC073::H2B-YFP* reporter lines, the last nucleated cell was defined as position 19 in each root corresponding to the average enucleation position obtained from live imaging. This way, the cells in similar developmental stages were placed closer together and thus the variance was reduced.

### Histological analyses and confocal microscopy

Cross-sections of hard resin-embedded Arabidopsis roots were obtained as described (*2*), stained with toluidine blue and imaged on Zeiss Axio Imager microscope equipped with 64 MP colour SPOT Flex high resolution camera. DAPI staining was performed as described (*3*). Confocal imaging was carried out on a Leica TCS SP8 equipped with 405 nm (DAPI stained samples), 442 nm (mTurq), 488 nm (GFP), 514 nm (YFP) and 561 nm (RFP and propium iodide stained samples) lasers. Live imaging was performed as described (*5*) on a Leica SP8 inverted scanning confocal microscope that uses solid-state 514 nm and 552 nm lasers for imaging YFP and mCherry respectively.

### Whole mount in situ hybridization

Target mRNA sequences were PCR amplified using whole seedling cDNA library of the Col-0 as a template (primer sequences are listed in Table S19). PCR fragments were cloned into pCR-Blunt II-TOPO vector (Invitrogen) that encodes both T7 and SP6 promoter priming sites. XbaI and HindIII restriction enzymes were used to linearize the APL and NEN4, NAC86 vectors, respectively, leaving only one of the promoters intact. In-vitro transcription reactions were performed using DIG RNA Labelling Kit (Roche). Hybridization and detection were carried out as described (*23*).

### Chromatin immunoprecipitation and qPCR

1/3 of 7-day-old roots was cut and 1.5g of root materials was collected in chilled cross-linking solution (1% formaldehyde and 5 mM EDTA in 1× PBS). Cross linking was done by applying vacuum for 30s, five times at room temperature. ChIP experiment was performed according to the published protocol (*36*) except for the DNA shearing step. Shearing was done in the BioRuptor, 30s on/off at 10 cycles. Primers used for the real-time PCR are listed in Table S19. Real-time PCR conditions were as follows: Initial denaturation 50°C 2 min followed by 95°C 10 min. Amplification was done with 45 cycles of 95°C 15 s, 60°C 1 min, and 72°C 15s. Reaction mix with SYBR was prepared as described in the manufacturer’s instruction (ROCHE). 3 independent biological replicates were used to calculate the fold-change with the BSA control set to 1.

### PLT2 induction experiment

To investigate PSE-abundant PLT2 targets, *pCALS7::H2B-RFP* was crossed with *pNAC86::XVE>>PLT2-3xYFP*. For the induction of PLT2, 5-day-old seedlings grown on the standard ½ media were transferred onto 10 μM 17-b-estradiol containing ½ MS media for 6 hours. In short, roots were cut and collected in the protoplasting solution containing the same volume of estradiol and DMSO respectively, and incubated for 12 min with vigorous swirling with forceps. After additional 50 min of incubation in the same solution, protoplasts were precipitated and re-suspended for the following cell sorting.

### Inducible and tissue specific manipulation of PLT2 expression with CRISPR-Cas9 technology

We used *plt12;pPLT2::PLT2-3xYFP*, partially complementing *plt1;2* double mutant phenotype, as a genetic background to CRISPR PLT2. Estradiol inducible *pPEAR1::XVE* promoter was used to drive CAS9p-t35S and four guide RNAs were designed for targeting *PLT2* as described (*20*).

### Fluorescence recovery after photobleaching to study precise expression domains

To precisely determine the expression domain of *pGIS::3xYFP*, Fluorescence recovery after photobleaching (FRAP) was used to abolish the fluorescence in all phloem cells and images of new translated proteins were taken after 90-120min of recovery. FRAP was done on a Leica TCS SP8 using the FRAP software module with activated FRAP booster. The laser lines 458 nm, 476 nm, 488 nm, 496 nm and 514 nm were simultaneously used at 70% laser power to bleach the phloem with a defined ROI at 256 × 256 pixel with 600 Hz scanning speed and with a x63 lens. 20-40 repetitions of the bleaching were applied for nearly complete bleaching. After recovery, regular confocal microscopy images were taken and the fluorescence quantified using the Fiji software package of ImageJ (*37*). The background fluorescence in the ground tissues was measured and subtracted from the measured fluorescence in the PSE cells. For each observed root, the mean fluorescence in the brightest cell was set to 1 and the others cell as a ratio of that.

### Cell behavior analysis

Long-term live imaging was performed with a nuclear localised reporter line (*pPEAR1::H2B-YFP pCALS7::H2B-YFP)* expressed in every cell of the PSE lineage. This reporter line was obtained by crossing *pPEAR1::H2B-YFP*, expressed strongly in positions 1-15^th^, with *pWOX5-mCherry/PET111-GFP*. F1 generation was then crossed to *pCALS7::H2B-YFP*, whose expression is predominant in the last 6 cells before enucleation (*3*). The imaging chamber was prepared according to the published method (*5*) with 1.5% agarose in the media.

### Random forest model for gene regulatory network inference

To model gene regulatory connections, we first selected the 15% most variable genes among the 758 cells using the genevarfilter function in Matlab (‘Percentile’, 85), leaving 4,924 genes for model inference. We modeled gene regulation using a Random Forest machine learning approach on the 758 single-cell expression profiles and the 4,924 highly variable genes, which included 208 transcription factors (TFs). In general, the Random Forest model allows for non-linear dependencies of target genes on causal transcription factors. Each single-cell expression profile is treated as a steady-state condition, allowing the model to learn a function that maps expression values of TFs to the expression value of each target gene. In the Random Forest approach, the TF-to-target association is described with a “score” that reflects the contribution of the TF to the expression of its target according to the model.

To address drop-out effects and other noise in single-cell data in the pre-processing stage, we merged the expression of consecutive cells to generate pseudo-cells using the following procedure to optimize the “bin size” (number of consecutive merged cells): we subdivided the 758 single-cell expression profiles into varying bin sizes, taking the median of the expression value of each gene in each bin or pseudo-cell as the value of that pseudo-cell. The Random Forest approach uses bootstrap aggregation, where each new tree is trained on a bootstrap sample of the training data. The remaining out-of-bag error is estimated as the average error for each training data point *pi* evaluated on predictions from trees that do not include *pi* in their corresponding bootstrap sample. For the dataset, the optimal bin size that minimized the out-of-bag error was 12 cells, providing our steady-state inference model a total of 64 pseudo-cells.

Finally, the Random Forest model ranks TFs based on their influence (score) on target gene expression, generating a predicted gene regulatory network (GRN) based on TF causality. To refine these TF–target predictions, we retained the top-10 highest scoring transcription factors for each gene target, resulting in 49,240 (TF-to-target) edges. The code is available at https://github.com/jacirrone/OutPredict. TFs were then ranked by their number of targets to derive the ranked list of the most important TFs (Table S14).

### Correlation analysis of the model in pseudotime

TF targets were classified into positive vs. repressive downstream sets in both the Random Forest model: Pearson correlation between each TFs and individual targets was used to determine regulatory effects (negative correlation, r<0, was classified as repressive regulation, and r>0 was classified as positive regulation, Table S14). We evaluated significant overlap between all pairwise positive and negative regulatory sets for each transcription factor (seesaw model) using the Fisher Exact test in Matlab. For the heatmap in Figure 3K, the binary output of the Fisher Exact test (p<0.05=1, p>0.05=0) was multiplied by the fraction of overlap between the two TF target sets (Table S15). Similarly, the overlap of up and down regulated targets in RNA sequencing upon ectopic overexpression of PLT2, PEAR1/PEAR2 (*23*) and NAC45 (*3*), ZAT14 or ZAT14-like was evaluated by systematically testing the pairwise intersection between the up- and down-regulated sets for each TF in the in vivo overexpression results. The criteria for identifying the up- and down-regulated sets (Set 1, Set 2) was an adjusted p value of 0.05 accounting for multiple testing and a two-fold change in expression over control either up- or down-regulated. The intersection of the two sets was tested for significant overlap using the Fisher Exact test, with a pval cutoff at 0.05. (Table S16). In the Fisher Exact tests, the overlap of Set 1 with all eligible transcripts was taken as outcome 1 (background intersection), while the overlap of Set 1 with Set 2 was taken as outcome 2 (test intersection).

### Live imaging of the late differentiating cells

*pNAC86::H2B-YFP* and *pNEN4::H2B-YFP* lines were crossed with *pWOX5-mCherry*. F1 seeds were grown vertically for 3 days on ½ MS plate and subsequently transferred to the imaging chambers described above to record movies S3-S12. Movies were analyzed to determine the time from the onset of H2B-YFP expression until the time given cell enucleates (fig. S1I).

## Supplementary Text

### Fluorescence-activated cell sorting

The destruction box (DBOX) sequence of *AtCYCB1;1* was fused to yellow fluorescent protein (YFP) to readily shut down YFP expression in dividing cells beyond promoter activity (*37*). Expression of this fusion protein using *PEAR1* and *CVP2* promoters resulted in predominant expression in the 2-6th and 3-9th cells, respectively (fig. S2A, B). However, these reporter constructs showed a cell-cycle biased expression pattern, in that the signal disappears right after cell division and comes back first in the cytoplasm and then to nucleus right before division (Movie S2). *pPEAR1* driven *3YFP* expression spans 10 cells from +2 position. For three other reporter lines, we combined two phloem reporters to limit the sorting population. For instance, *pNAC057::2GFP* was combined with *pCALS7::H2B-RFP*, which restricts the only GFP positive cell population to cells 4-14. (fig. S2A, B). The expression of *pCALS7::H2B-YFP* is observed in 7 cells prior to the enucleation, whereas NAC073 promoter is active in a narrow domain consisting of 4 cells before enucleation. However, YFP fluorescence could be found also in phloem pole pericycle cells neighboring enucleated PSE in both lines (fig. S2B). We’ve combined these with *pS32::erRFP*, a reporter broader in the stem cell niche and exclusive in differentiating PSE, creating a refined overlapping expression in PSEs.

### Definition of seven domains

The stem cell (I) is defined as a first cell of the PSE lineage, positioned adjacent to the quiescent center (dark gray in panel A of figure 1) and divides in a self-renewing manner. In each of the two movies, only one division of the stem cell was observed. Based on this, we conclude that division of the stem cell likely does not happen more often than every 60h.

Transit amplifying cells (II), positions 2-9, are anticlinally (amplification of the cells within the PSE lineage) or periclinally (contributing to MSE and procambium) dividing cells. Transit amplifying cells undergo multiple divisions over a minimum of 58h before they enter the “Transition” zone (III), positions 8-11, where they divide for the last time after which both daughter cells initiate process of differentiation. As the “transition” is defined by a single event, i.e. last cell division, the time spent in this zone cannot be measured. Division of any TA cell gives rise to a rootward and a shootward daughter. The time cells spend in the domain II was calculated for the faster progressing, shootward daughter cells. This time includes also, on average, half of the cells from the transition zone.

Process of differentiation, measured here from the last cell division of transitioning cells till enucleation, took on average ~20h). Because differentiating cells do not proliferate, the time they spend in each position depends on the frequency of enucleation which “removes” one cell at the time from the end of the differentiating PSE lineage. In the two recorded movies enucleation took place, on average, every 2 hours. Following this, the “Enucleating cell” in position 19, domain (VII), spends 2h in this position. This value was the basis to calculate the timing of the other differentiation domains. The time cells spend in the “Early differentiation” domain (IV), positions 10-15, has been estimated to take 12h, “Late differentiation” (V) positions 16-17, 4h, and “Very late differentiation” (VI) positions 18-19, 4h.

## Acknowledgements

We thank Yuki Kondo, Philip Benfey and Ottoline Leyser for providing seeds of *pNAC57::2GFP*, *pSHR::erGFP* and *pPIN4::PIN4-GFP* reporters, respectively and the staff of the Flow Cytometry Core Facility at CIMR for their technical support with cell sorting. We are thankful to Raymond Wightman and Gareth Evans for technical support with microscopy experiments. This work was supported by Finnish CoE in Molecular Biology of Primary Producers (Academy of Finland CoE program 2014-2019) decision #271832, the Gatsby Foundation (GAT3395/PR3)), University of Helsinki (award 799992091) and the ERC Advanced Investigator Grant SYMDEV (No. 323052) to Y.H., a NSF-BBSRC MCSB grant (1517058) to R.S. and Y.H., a NSF CAREER MCB grant (1453130) to R.S., a NIH grant (GM078279) to K.D.B., a NFS IOS grant (1934388) to K.D.B and D.S., a MRC Clinical Research Infrastructure award (MR/M008975/1) and Core funding from the Wellcome and MRC to the Cambridge Stem Cell Institute to B.G., Academy of Finland grants (266431 and 307335) to A.P.M. and X.W., an NWO Horizon grant (050-71-054) to R.H., ERC Starting Grant (TORPEDO; 714055) and the Research Foundation - Flanders (FWO; Odysseus II G0D0515N) to B.D.R., a Gatsby Foundation CDF grant to S.E.A., a BOF postdoctoral fellowship from Ghent University to J.R.W., a Wallenberg Academy Fellowship (KAW 2016.0274) to C.M., a Wellcome Strategic Award (105031/D/14/Z) to F.H., a JSPS Research Fellowship for Young Scientists and JSPS KAKENHI grant (JP16J00131) to K.T., and The Finnish Academy of Science to J.H.

## Authors contributions

P.R., J.H., B.B., W.W.Y.L., R.U. performed experiments and collected data., P.R., J.H., B.B., K.T., M.A.d.L.B., F.H., J.C., H.T., K.V., J.W., B.D.R., S.E.A. and K.D.B. analyzed data., J.C. and X.W. developed methodology., P.R., J.H., B.B., K.T., M.A.d.L.B., J.C., A.P.M., R.S., K.D.B. and Y.H. conceptualized and designed the study, P.R., J.H., B.B., K.D.B. and Y.H. wrote the manuscript. All authors read, edited and discussed the manuscript.

## Data and materials availability

All data are available in the supplementary material and raw RNA sequencing data for pooled (bulk sorted) PSE cells, single PSE cells and overexpression and mutant profiling is deposited at GEO-NCBI (GEO submissions GSE142259, GSE140778, GSE140977, respectively).

## Supplementary tables

All Supplementary Tables can be found in the different work sheets in the Supplementary_Tables.xlsx file.

Table S1: List of fluorescence reporter and mutant lines of *A. thaliana* analyzed in this publication. Genetic background, introduced transgenic construct, abbreviation and origin are given.

Table S2: List of temporal specific genes. These were identified by comparing the transcriptomes in this domain to those of all others in pairwise comparisons. The highest qval of all these comparisons was then used for further selection (only genes with qval <0.05 are listed). Genes specific to the nested PLT1-like and NEN4-like domains can also be part of domain [a] or domain [d], respectively.

Table S3: List of PLT targets after ectopic expression in late PSE cells. The expression in each replicate (RPKM), logarithm of the fold change (logFC, between EST/induced and DMSO/non-induced), p values (PValue), false discovery rates (FDR) are given. Upregulated genes that have GO term annotations as “cell cycle” are indicated as well as here detected specificity to M or S phase of those genes.

Table S4: List of 925 phloem enriched genes based on Shannon entropy analysis of bulk sorted cells. The gene IDs, names and description are given. RPKM values for all used replicates, including the published data run through the same bioinformatics pipeline are given (Columns D-BK). Statistics of the Shannon entropy calculations can be found in Columns BL-BS.

Table S5: List of 1192 SE enriched genes based on cluster comparisons of 272 *pPEAR1Δ::erVENUS* single cells from the PSE, MSE and procambial lineage. For further description of the applied comparisons, see M&M.

Table S6: List of the PSE enriched genes that form the overlap between Table S4 and Table S5.

Table S7: Average expression per cluster of 467 out of 542 phloem enriched genes (Table S6) that were detected in published root expression atlas (Wendrich et al., submitted). These genes are not broadly expressed throughout the tissue types. While many genes show either PSE or phloem specific expression, some show specific expression in other cell types. Several genes confirmed to be expressed during late PSE cells such as *NAC010, NAC73, ZAT14, RMT2* and *NEN4* appear not to be expressed in the sieve element cluster. This could potentially indicate the underrepresentation of late PSE cells within the sieve element cluster (<30 cells).

Table S8: Core predictions of the random forest GRN. The top10 most important transcription factors for each gene were selected, resulting in 49240 edges in the model (ranked by importance). The gene ID for transcription factor (TF) and the predicted target (Target), the importance of this edge (Importance), the correlation (Correlation) and the regulation based on the correlation is given (Correlation sign). These core edges of the model were in turn used to determine the most important transcription factors by the number of their target genes. The gene ID for each transcription factor (gene), number of target genes (target #), combined importance of these edges (target importance) and a short annotation for each transcription factor (annotation) is given. The model was tested for enrichment of targets determined in available overexpression or mutant data sets. The results of this enrichment test also given.

Table S9: List of differentially expressed genes in the pear sextuple root meristem. Read counts, log2 fold change, pval and padj are given. Only significantly differentially expressed genes (padj<0.05) are shown.

Table S10: List of differentially expressed genes after 10h induction of *pWOL::XVE>>ZAT14*. Gene IDs, mean read counts of 3 replicates for 10h induction and DMSO control (Z10 and Z_DMSO), log2foldchange, pvalue, padjust, gene loci and gene names are given for all significantdifferentially expressed genes (padj<0.05).

Table S11: List of differentially expressed genes after 14h induction of *pWOL::XVE>>ZAT14*. Gene IDs, mean read counts of 3 replicates for 14h induction and DMSO control (Z14 and Z_DMSO), log2foldchange, pvalue, padjust, gene loci and gene names are given for all significant differentially expressed genes (padj<0.05).

Table S12: List of differentially expressed genes after 10h induction of *pWOL::XVE>>ZAT14L*. Gene IDs, mean read counts of 3 replicates for 10h induction and DMSO control (L10 and L_DMSO), log2foldchange, pvalue, padjust, gene loci and gene names are given for all significant differentially expressed genes (padj<0.05)

Table S13: List of differentially expressed genes after 14h induction of *pWOL::XVE>>ZAT14L*. Gene IDs, mean read counts of 3 replicates for 10h induction and DMSO control (L14 and L_DMSO), log2foldchange, pvalue, padjust, gene loci and gene names are given for all significant differentially expressed genes (padj<0.05)

Table S14: PANTHER Gene ontology analysis of ZAT14 and/or ZAT14L targets (14h induction). Details such as test type and correction are given alongside the expected and detected number of genes for each GO category, the fold enrichment and p value.

Table S15: Significant overlap of up and down regulated, predicted targets for the Top20 transcription factors in the model, respectively.

Table S16: Summary of statistical tests for overlapping gene sets of downstream targets in over-expression data sets. Overrepresentation ratio and pval (Fisher test) are given as well as genes that are both up and down regulated by early or late transcription factors. PLT2 and ZAT14L were included as those are also important regulators in this study with very specific early or late expression and overexpression target data was available.

Table S17: List of single cell transcriptomes used in this study. Cell identifiers (ID), reporter line used to isolate the cell (sample), number of total reads, unmapped reads, reads mapped to a single locus and reads mapped to multiple loci, percent of reads mapped to single locus, reads mapped to multiple loci and generally mapped to A. thaliana genome, average library size (lib_size_avr), Cluster number when all cells passing quality thresholds were clustered (Cluster_all_cells), cluster number when only 758 PSE were clustered (Cluster_PSE) and assigned pseudotime for PSE cells are given (Pseudotime_PSE).

Table S18: Live imaging quantification data. The time between onset of expression and enucleation in *pNAC86::H2B-YFP* and *pNEN4::H2B-YFP* line and between enucleation was quantified. For the expression length, the root and corresponding movie, the observed phloem pole (pole), movie frame with onset and enucleation and the difference (frame gene ON, frame enucleation, frames gene expression, respectively), the time of expression before enucleation in hours (time cell enucleation [h]) and the corresponding movies are given. The enucleation quantification is based only on movie S1 and movie S2 and their analysis in Fig S1A, E. Time between consecutive enucleation events and the corresponding movie are given.

Table S19: List of oligonucleotide sequences used in this study. Primer names, including gene names and positions and amplified fragment sizes are given together with the sequence.

## Supplementary Movies

Movie S1: PSE cell division behavior movie 1. Cell divisions in the PSE cell file were quantified and are shown in Fig S1A using the *pPEAR1::H2B-YFP pCALS7::H2B-YFP pWOX5-mCherry* reporter line to follow the nuclei (YFP) and track the root (mCherry).

Movie S2: PSE cell division behavior movie 2. Cell divisions in the PSE cell file were quantified and are shown in Fig S1E using the pPEAR1::H2B-YFP pCALS7::H2B-YFP pWOX5-mCherry reporter line to follow the nuclei (YFP) and track the root (mCherry).

Movie S3: pNAC86::H2B-YFP_movie1. Duration of promoter activity of NAC86 (YFP positive nuclei) was quantified in this movie and is shown in Fig S1I and Table S18.

Movie S4: pNAC86::H2B-YFP_movie2. Duration of promoter activity of NAC86 (YFP positive nuclei) was quantified in this movie and is shown in Fig S1I and Table S18.

Movie S5: pNAC86::H2B-YFP_movie3. Duration of promoter activity of NAC86 (YFP positive nuclei) was quantified in this movie and is shown in Fig S1I and Table S18.

Movie S6: pNEN4::H2B-YFP_movie1. Duration of promoter activity of NEN4 (YFP positive nuclei) was quantified in this movie and is shown in Fig S1I and Table S18.

Movie S7: pNEN4::H2B-YFP_movie2. Duration of promoter activity of NEN4 (YFP positive nuclei) was quantified in this movie and is shown in Fig S1I and Table S18.

Movie S8: pNEN4::H2B-YFP_movie3. Duration of promoter activity of NEN4 (YFP positive nuclei) was quantified in this movie and is shown in Fig S1I and Table S18.

Movie S9: pNEN4::H2B-YFP_movie4. Duration of promoter activity of NEN4 (YFP positive nuclei) was quantified in this movie and is shown in Fig S1I and Table S18.

Movie S10: pNEN4::H2B-YFP_movie5. Duration of promoter activity of NEN4 (YFP positive nuclei) was quantified in this movie and is shown in Fig S1I and Table S18.

Movie S11: pNEN4::H2B-YFP_movie6. Duration of promoter activity of NEN4 (YFP positive nuclei) was quantified in this movie and is shown in Fig S1I and Table S18.

Movie S12: pNEN4::H2B-YFP_movie7. Duration of promoter activity of NEN4 (YFP positive nuclei) was quantified in this movie and is shown in Fig S1I and Table S18.

---

**Supplementary figure 1.**
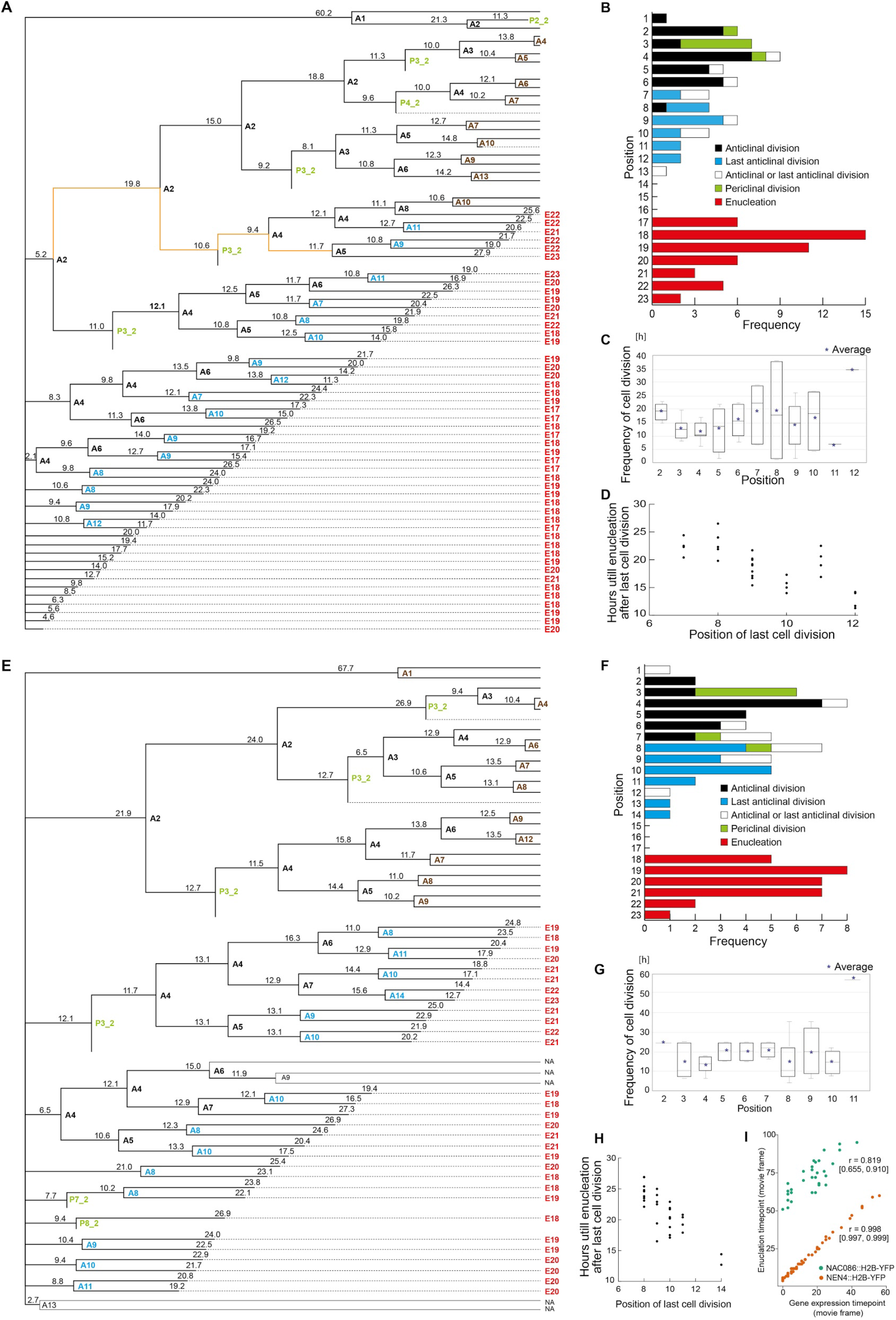
Live imaging of PSE development. A) and E) Cell division trees extracted from Supplementary movies 1 and 2, respectively. Time between cell divisions is indicated above the line connecting two nodes that demark type (A=anticlinal, P=periclinal) and position of the cell at the time of division. Cell division annotation in black describes anticlinal divisions in Transit Amplifying zone; in blue, last anticlinal divisions (Transition zone); in brown, last divisions observed in the movie where fate of the daughter cells can’t be determined; in green, periclinal divisions (no first periclinal division has been observed). Annotation in red informs about the position of enucleation. Division path in orange depicts shortest passage through transit amplifying cells observed. B) and F) Histograms of cellular events from movies 1 and 2, respectively. C) and G) Time between cell divisions in different positions of PSE cell lineage. Dots indicate the average. D) and H) The time from last cell division till enucleation plotted for different cell positions. I) Pearson’s correlation (r) between gene expression activation time point and the time point this cell enucleated. 95% confidence interval is provided in square brackets. The confidence interval is wider for *NAC086*, reflecting a higher variation in the interval between the time the gene expression is activated and the time the cell enucleates.

**Supplementary figure 2.**
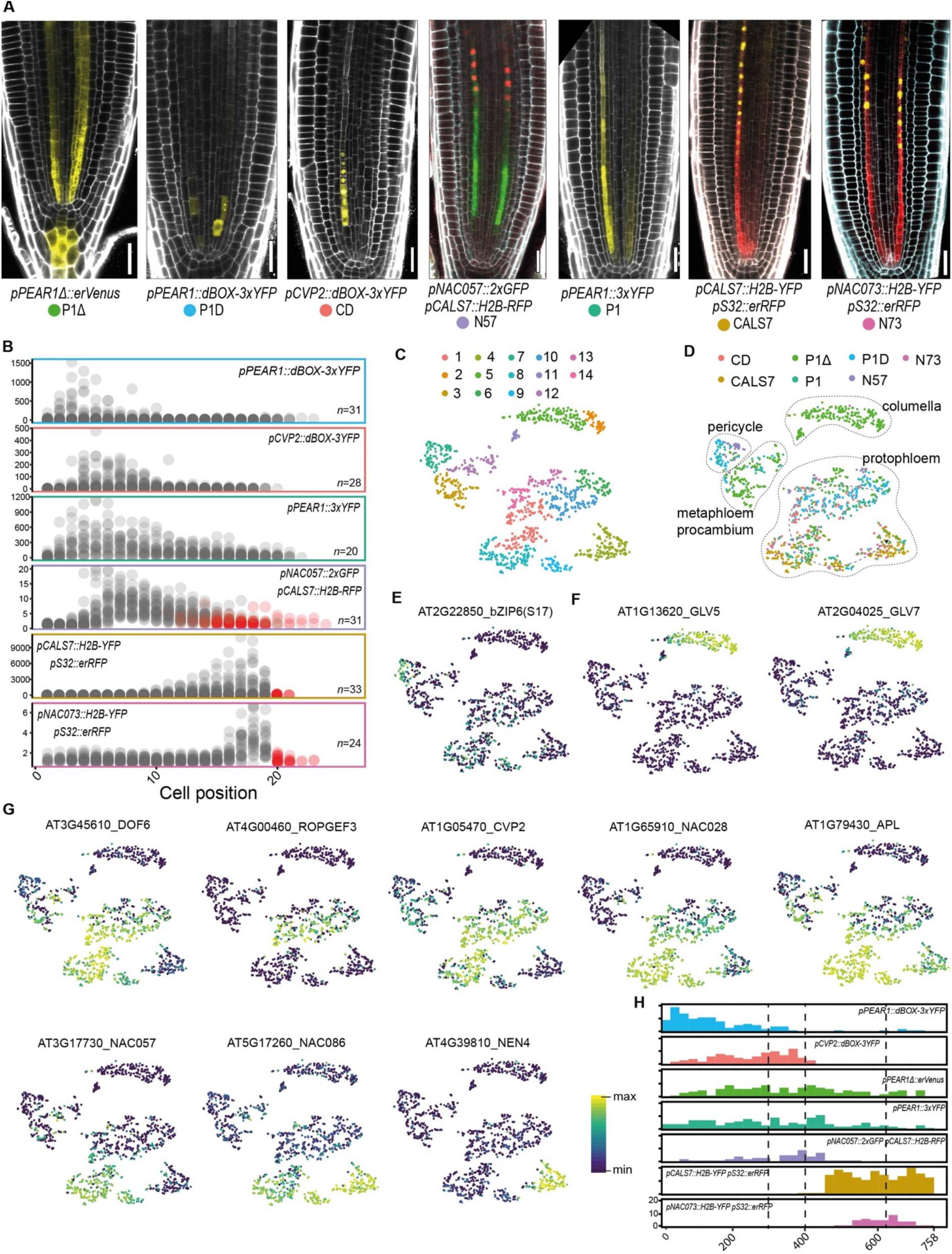
Sorting of phloem specific cells. A) Reporter lines used for protoplasting and FACS based isolation of cells. Modified *PEAR1* promoter (P1Δ) shows ectopic expression in columella reflected in additional columella cluster in (D, F). Scale bars, 25 μm. B) Fluorescent signal intensity distribution in the phloem reporter lines from (A). Averaged and background-normalised fluorescence per PSE cells is given from the stem cell to the enucleating cell. Red dots indicate cells with RFP fluorescent nucleus (*pNAC057::2xGFP pCALS7::H2B-RFP*) or that are enucleated (*pCALS7::H2B-YFP* and *pNAC073::H2B-YFP*). C) t-SNE plot reveals 14 different clusters of the analysed cells. D) t-SNE plot with transcriptomes color-coded based on the reporter line they originate from. Cluster annotation based on the expression of known, tissue-specific genes. E) Expression of pericycle specific and (F) columella specific genes indicates cluster identities. F) (*above*) G) Identification of PSE cluster(s) based on the expression of sieve element specific genes. Subsequent t-SNE plots indicate a developmental gradient within PSE cluster(s). H) Histogram of cell distribution along pseudotime of 758 ordered PSE transcriptomes based on the reporter line used for isolation. Correlation drop points are indicated with dashed lines.

**Supplementary figure 3.**
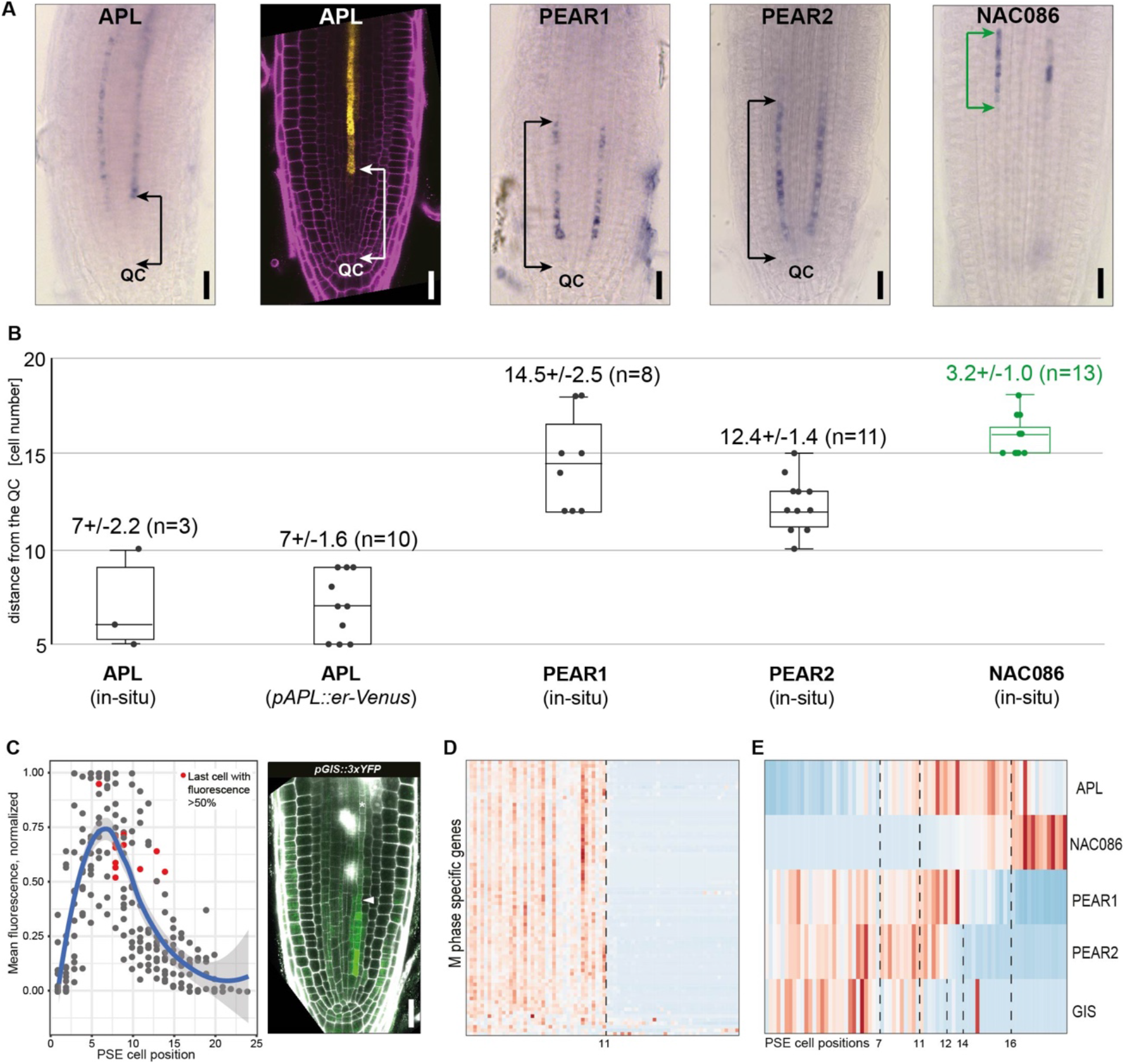
Mapping of the single cell transcriptome onto 19 cells of the PSE cell lineage. A) Representative images of in situ RNA hybridization and reporter line indicating the onset or termination of expression of PSE specific genes. The quantified domains are indicated with arrows. Scale bars, 25 μm. B) Quantification of domains indicated in panel (A) representing onset of *APL* and *NAC086* and termination of *PEAR1* and *PEAR2* expression. Expression of *NAC086* (in green) terminates upon cell enucleation at position 19. C) Quantification of relative fluorescence per PSE cell position of *pGIS::3xYFP* lines in FRAP experiments (n=12 phloem poles). Representative image of *pGIS::3xYFP* after FRAP recovery. Arrowhead points at the last cell with fluorescent level above 50%. Scale bar, 25 μm. D) Gene expression heatmap of genes specifically expressed during M phase of the cell cycle (Menges et al., 2005). The end of expression can be linked to the position 11 in the live imaging data of cell division in figure S1. E) Heatmap showing expression of genes quantified in (A to C) in pseudotime ordered PSE transcriptomes. Approximate start or end points of gene expression are indicated with dashed lines, linking quantified positions in (B, C, D) to pseudotime positions.

**Supplementary figure 4.**
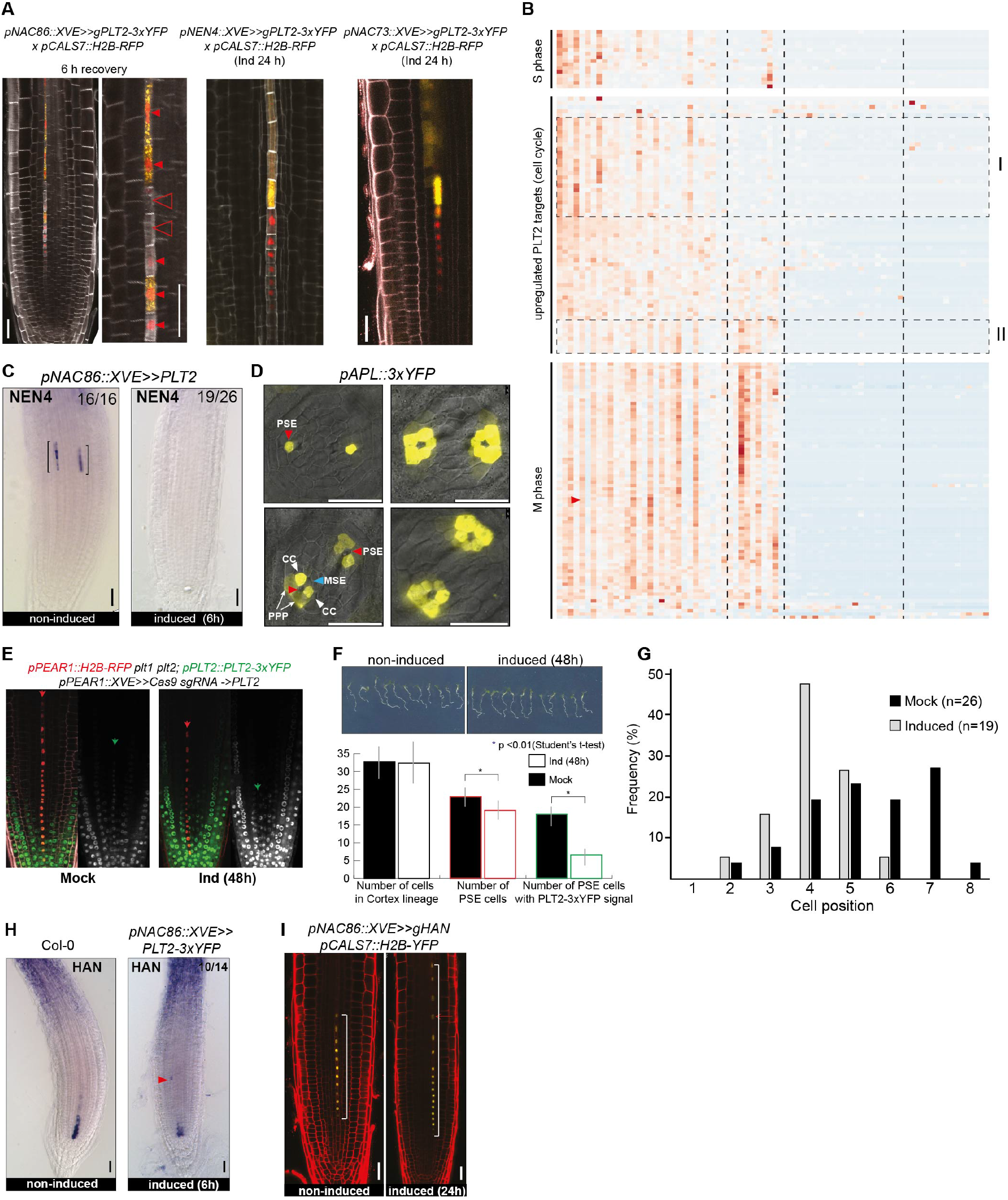
PLT2 inhibits expression of late PSE genes and the key regulator APL. A) The effect of ectopic expression of *PLT2* under p*NAC086* is reversible as indicated by the enucleation of PLT2-free PSE cells after recovery. During very late differentiation of phloem (domain VI) expression of *PLT2* under *pNAC073* and *pNEN4* does not delay enucleation, neither promotes cell division. B) Gene expression heatmap highlighting oscillatory patterns of a subset of PLT2 upregulated cell cycle related targets (clusters I and II), coinciding with expression of either S or M phase specific genes. C) In situ hybridization of *NEN4* before and 6h after ectopic expression of PLT2-3xYFP. Brackets indicate in situ signal. D) Expression of *APL* in the PSE before enucleation and its expression in phloem pole pericycle (PPP), companion cells (CC) and metaphloem sieve element (MSE) after PSE enucleation. E) Premature enucleation of phloem cells, monitored by *pPEAR1::H2B-RFP*, after 48h induction of PSE specific CRISPR construct targeting *PLT2* (including complementing construct) in *plt1 plt2 pPLT2::PLT2-3xYFP* background. Last nucleated PSE cell and last PSE cell expressing PLT2-3YFP are indicated with red and green arrows, respectively. F) No change to meristem size was observed 48h after induction of CRISPR construct. Number of PSE cells was monitored based on *pPEAR1::H2B-RFP*. G) Quantification of p*APL::erTurq* expression after induction of tissue specific CRISPR construct targeting *PLT2*. H) Native and ectopically activated expression of *HAN*. Arrowhead indicates ectopic expression of *HAN* promoted by ectopic expression of PLT2. I) Ectopic expression of HAN delays enucleation in the late domain of differentiating phloem. Square brackets indicate extended expression domain of *pCALS7::H2B-RFP*, a reporter used for monitoring enucleation. All scale bars, 25 μm.

**Supplementary figure 5.**
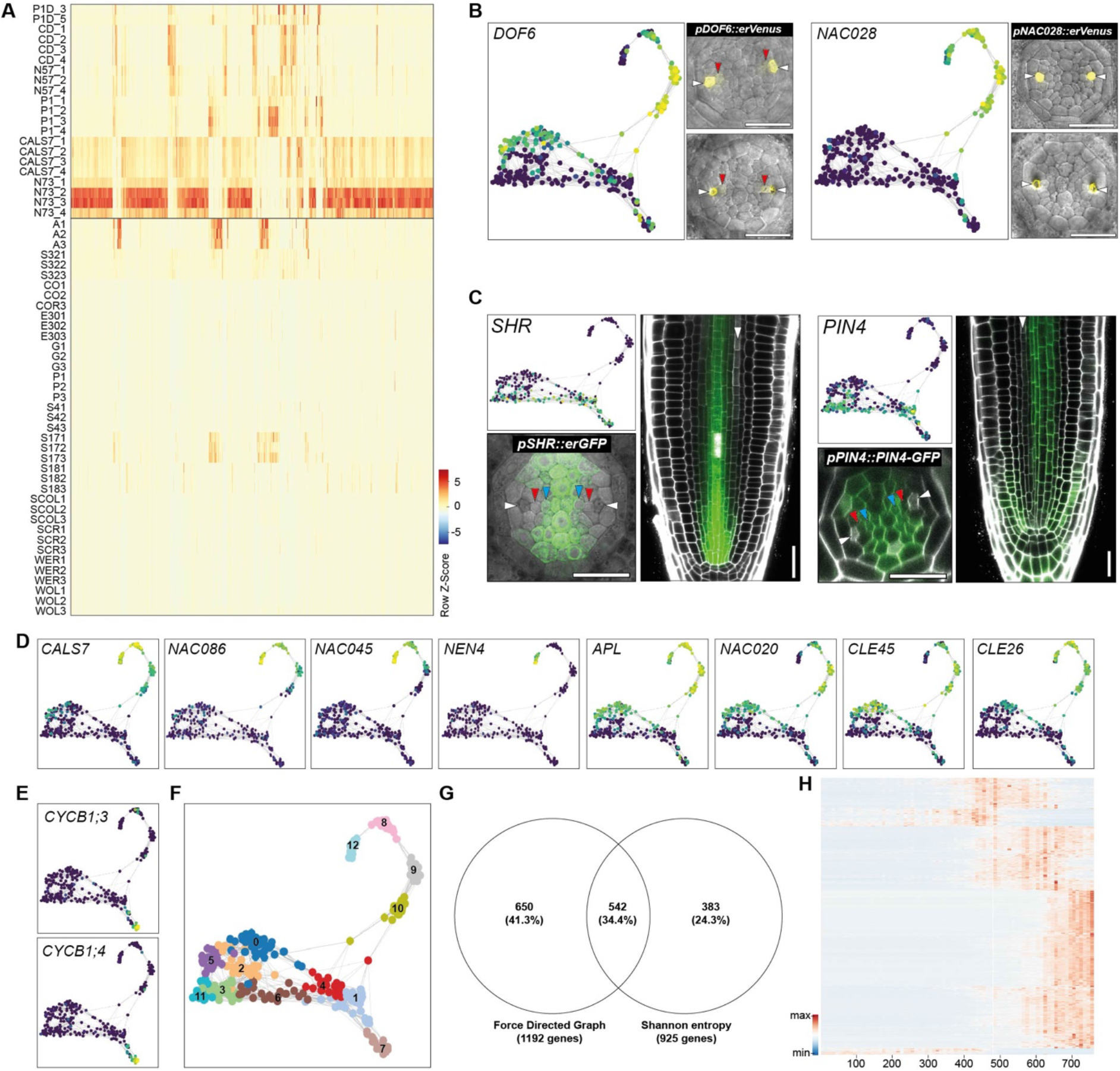
Identification of PSE enriched genes using bulk and individually sorted cells. A) Expression heatmap of 925 phloem enriched genes in the combined transcriptome data set of bulk sorted PSE domains and other cell types from Li et al., 2016. B) Force-directed graph of 272 single-cell transcriptomes obtained using the *pPEAR1Δ::erVENUS* reporter. Expression of known sieve element genes indicates protophloem and metaphloem sieve element trajectories. Vibratome cross sections of fluorescent reporter lines confirm SE specificity of both genes. C) Identification of procambial (“non-SE”) cell trajectory based on the expression pattern of *SHORT ROOT* (*SHR*) and *PIN-FORMED 4* (*PIN4*). Both genes are expressed broadly in the very early cells but quickly become excluded from the PSE and MSE cell files. D) Expression pattern of known sieve element specific genes in the force-directed graph analysis confirms cluster identity. E) Expression of M-phase specific cyclins indicates the separation of cluster 7 (F) is based on cell cycle phase. F) Distribution of 12 Louvain clusters of the single cell transcriptomes from cell sorting of *pPEAR1Δ::erVENUS* reporter. G) Venn diagram of phloem enriched (Shannon Entropy analysis of bulk sorted cells) and sieve element specific genes (based on the comparisons between Louvain clusters) shows the large overlap of 542 common genes. Expression heatmap of 542 SE enriched genes from (G) in pseudotime. SE enriched genes show predominant expression during late stages of PSE development.

**Supplementary figure 6.**
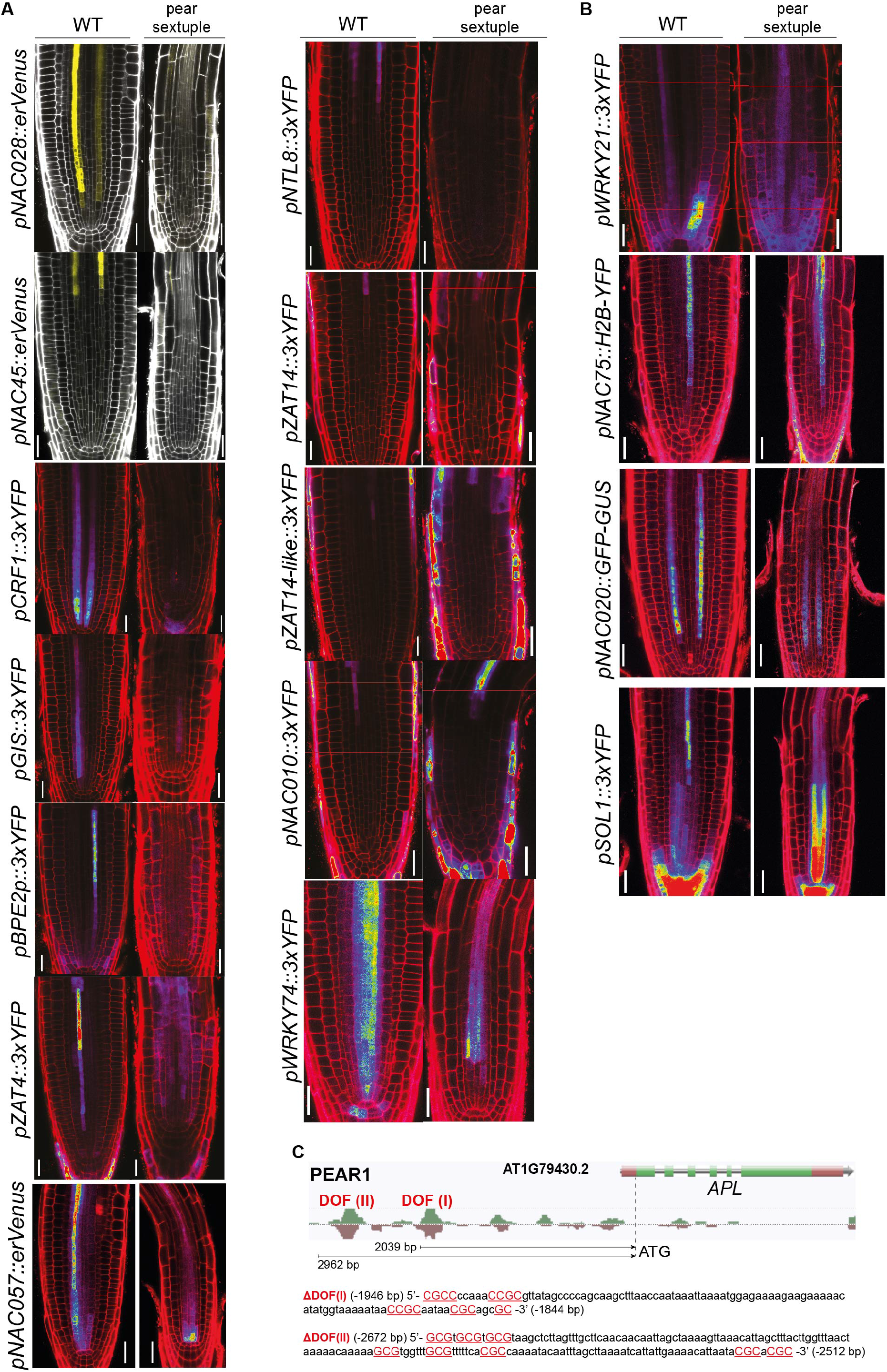
Expression pattern of PEAR-dependent and - independent, PSE enriched TFs. A) Known and newly identified PSE-enriched TFs whose expression during phloem development depends on PEARs. B) 4 of the tested promoters remained active in pear sextuple mutant, indicating that expression of these transcription factors does not depend on PEARs. Images were taken using the same settings, ensuring comparable intensities between genetic backgrounds. All scale bars, 25 μm. C) An image extracted from Plant Cistrome Database (http://neomorph.salk.edu/dev/pages/shhuang/dap_web/pages/index.php) showing PEAR1 binding to *APL* promoter. DOF binding sites of the two prominent peaks have been modified as listed below the image. Coordinates indicate position upstream of APL ORF.

**Supplementary figure 7.**
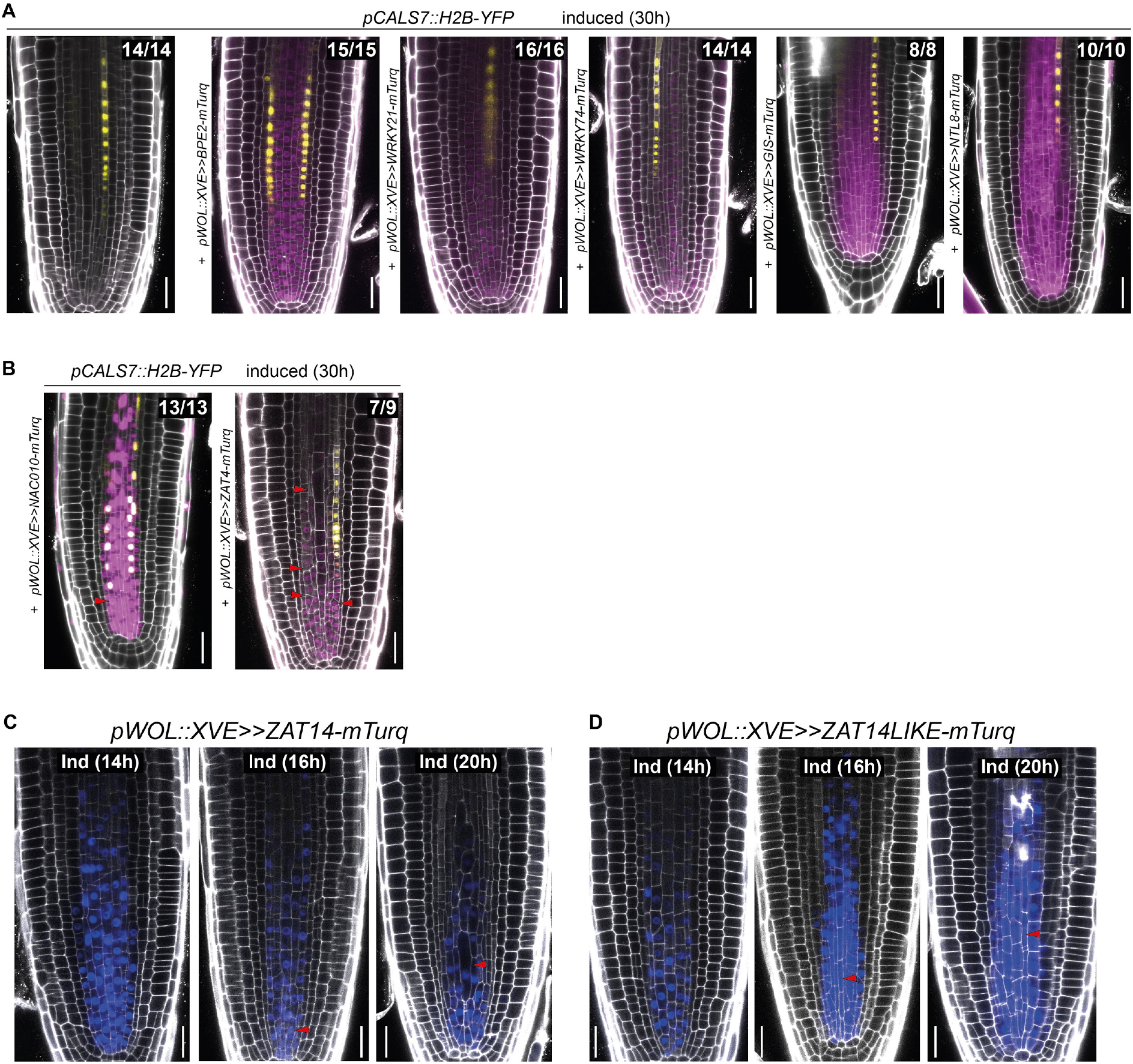
Ectopic overexpression studies of newly identified PSE enriched transcription factors. A) Overexpression of *BPE2*, *WRKY21*, *WRKY74, GIS* and *NTL8* under *pWOL::XVE* promoter, active broadly in the vascular tissue, did not result in any obvious phenotypes. Transgenic lines carry *pCALS7::H2B-YFP* to monitor phloem enucleation. B) Overexpression of *NAC010* and *ZAT4* resulted in shortening of the root meristem and defects in the cell shape, respectively, indicated by red arrowheads. C) and D) Extensive cell elongation and inhibition of cell division phenotypes in the lines overexpressing *ZAT14* and *ZAT14-like*. Red arrowheads indicate unusual cell expansion. First signs of the phenotype appear at 16h after induction. Lines were profiled (RNA-Seq of root meristems) at 10h and 14h after induction. Numbers indicate observations with similar expressions and phenotype. All scale bars, 25 μm.

**Supplementary figure 8.**
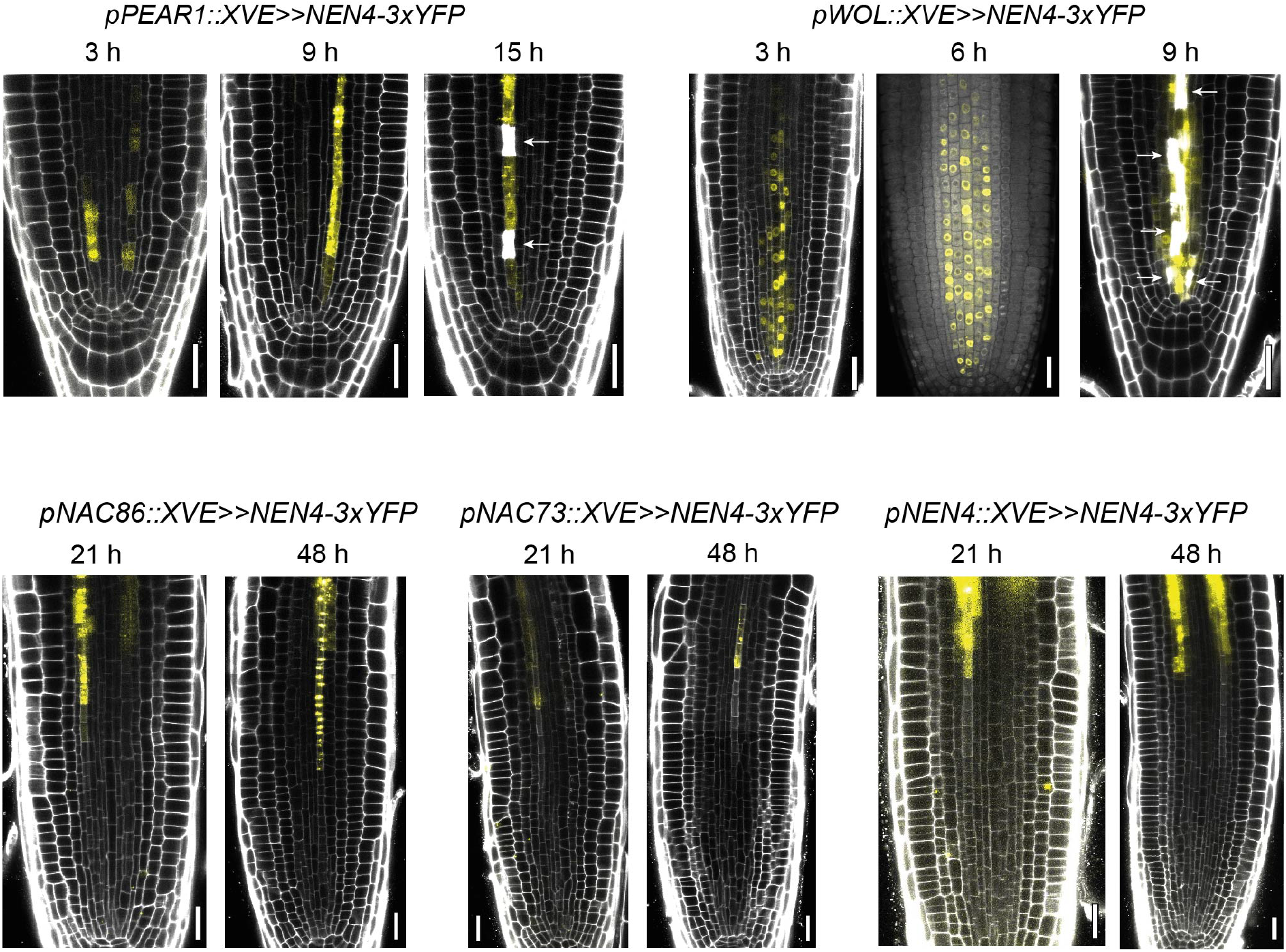
Ectopic expression of NEN4 reveals stage of fixed PSE cell identity. Premature expression of *NEN4* under PSE-specific *pPEAR1* or the broadly active *pWOL* promoters results in cell death after 15 or 9 hours, respectively. On the other hand, expression of *NEN4* under late PSE promoters (*pNAC86, pNAC73, pNEN4*) reveals resistance of cells close to maturation to NEN4-induced cell death. White arrows point to dead cells. All scale bars, 25 μm.

**Supplementary figure 9.**
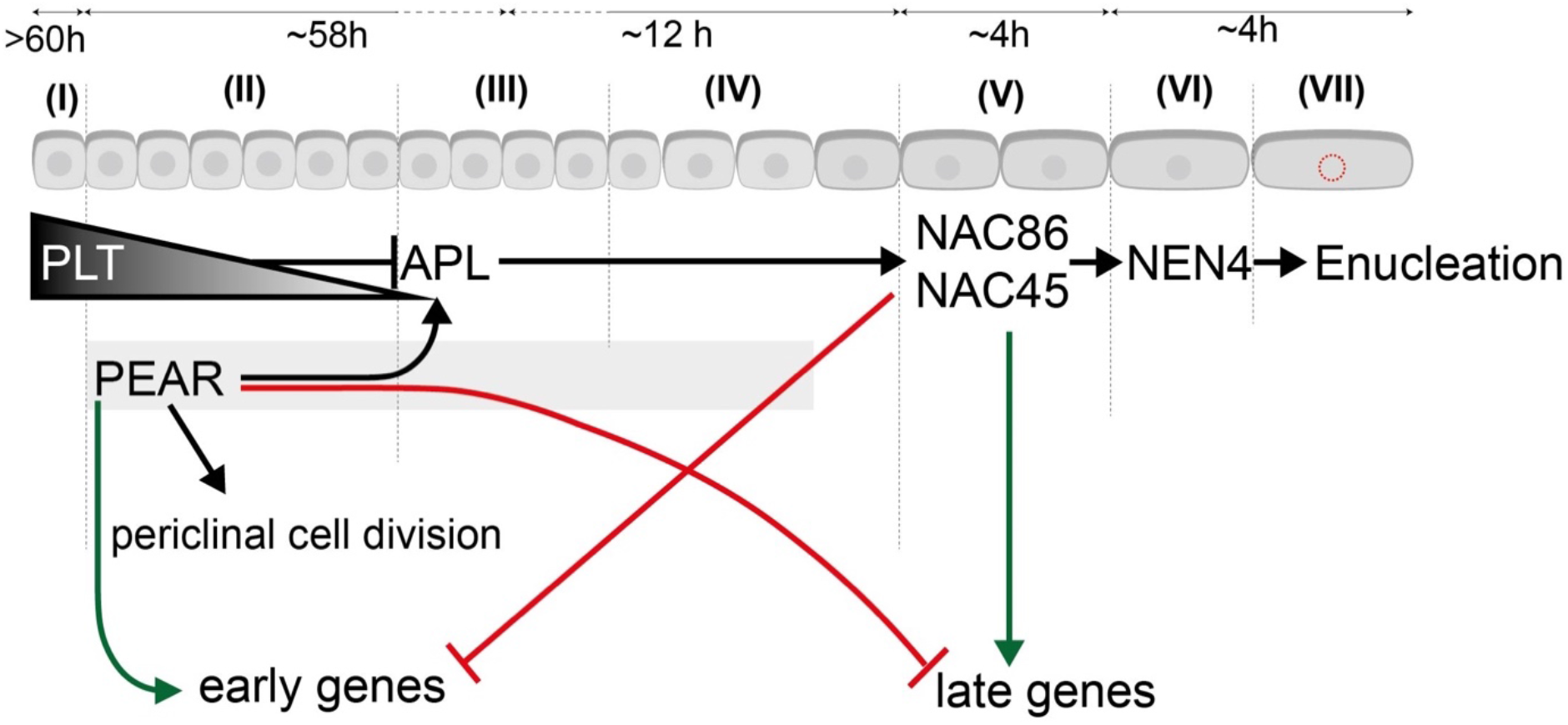
Gene regulation along PSE developmental trajectory. Schematic summarizes genetic interactions between major transcription factors orchestrating PSE development imposed on the PSE trajectory for which zonation and timing of the events has been determined in this study. The seesaw concept is indicated by the validated up- and down-regulation of early and late expressed genes by PEARs and NAC45/86, respectively and *vice versa*.

